# Transcriptomically-guided mesendoderm induction of human pluripotent stem cells using a systematically defined culture scheme

**DOI:** 10.1101/561068

**Authors:** Richard L Carpenedo, Sarah Y Kwon, R Matthew Tanner, Julien Yockell-Lelièvre, Chandarong Choey, Carole Doré, Mirabelle Ho, Duncan J Stewart, Theodore J Perkins, William L Stanford

## Abstract

Human pluripotent stem cells (hPSCs) are an essential cell source in tissue engineering, studies of development, and disease modeling. Efficient, broadly amenable protocols for rapid lineage induction of hPSCs are of great interest in the stem cell biology field. We describe a simple, robust method for differentiation of hPSCs into mesendoderm in defined conditions utilizing single-cell seeding (SCS) and BMP4 and Activin A (BA) treatment. Gene sets and gene ontology terms related to mesoderm and endoderm differentiation were enriched after 48 hours of BA treatment. BA treatment was readily incorporated into existing protocols for chondrogenic and endothelial progenitor cell differentiation. After prolonged differentiation in vitro or in vivo, BA pre-treatment resulted in higher mesoderm and endoderm levels at the expense of ectoderm formation. These data demonstrate that SCS with BA treatment is a powerful method for induction of mesendoderm that can be integrated into protocols for mesoderm and endoderm differentiation.

## Introduction

Pluripotent stem cells (PSCs) are a powerful tool in a variety of applications ranging from basic studies of development and disease to cell-based therapeutics and regenerative medicine applications (Evans and Kaufman, 1981; Martin, 1981; Reubinoff et al., 2000; Thomson et al., 1998). Human embryonic (hESCs) (Thomson et al., 1998) and induced pluripotent stem cells (iPSCs) (Takahashi et al., 2007; Yu et al., 2007) are two classes of PSCs that are particularly well-suited for modeling genetic diseases (Dimos et al., 2008; Park et al., 2008; Soldner et al., 2009) as well as serving as a renewable cell source for tissue engineering purposes (Tabar and Studer, 2014; Wu and Hochedlinger, 2011). For the potential of PSCs to be realized from basic science to clinical applications, efficient directed differentiation protocols to produce relevant cell types are required. While much work has been done in the area of inducing differentiation of PSCs to various types of somatic cells, methods for generating cells of interest that are simple, chemically defined, and can be adapted and optimized for many cell types are of great interest to a breadth of scientists, engineers, and clinicians.

During gastrulation, pluripotent epiblast cells migrate through the primitive streak, leading to formation of the mesoderm and definitive endoderm germ lineages. Morphogens secreted by surrounding embryonic tissues, such as BMP4, Activin A, Wnt, and FGFs, are responsible for orchestrating the complex and dynamic morphogenesis that results in germ layer formation (Tam and Loebel, 2007; Tam et al., 2006). While gastrulation has been studied extensively in the mouse, a number of differences exist between human and mouse embryos, including the shape of the epiblast, (flat in human, cup-shaped in the mouse (Solnica-Krezel, 2005)), the influence of the epiblast-chorionic ectoderm boundary on radial symmetry breaking (Sheng, 2015), and the spatio-temporal patterns of transcription factor expression in the blastocyst (Niakan and Eggan, 2013). Similarly, stem cells derived from human and mouse embryos display important distinctions, including the factors required for self-renewal as well as differentiation capacity (Ginis et al., 2004; Wei et al., 2005). Thus, human PSCs are a compelling model system to investigate human development, as human ESCs and iPSCs more accurately recapitulate aspects of human development than the mouse embryo or mouse PSCs.

Many cell types important for tissue engineering applications are mesendoderm derived, including chondrocytes, vascular cells, cardiomyocytes, hepatocytes, and pancreatic β-cells. A number of physical parameters influence hPSC differentiation to these cell types, including cell density, colony size, and dissociation method, as hPSCs are exquisitely sensitive to paracrine factors from neighboring cells (Davey and Zandstra, 2006) and can undergo apoptosis and display karyotypic abnormalities when passaged as single cells (Draper et al., 2004; Mitalipova et al., 2005). Advanced microengineering approaches have been used to control cell spacing and colony size, resulting in differentiation platforms amenable to induction of multiple lineages (Bauwens et al., 2008; Blin et al., 2018; Lee et al., 2009; Mohr et al., 2006; Nazareth et al., 2013; Peerani et al., 2007). While microfabricated systems can be beneficial for enhancing microenvironmental control over differentiating cells, they are not practical for many laboratories performing fundamental studies. Thus, a simple and broadly applicable platform for controlling microenvironmental conditions that can be utilized in laboratories with a range of specialties to induce differentiation of human PSCs to mesendoderm is required.

Here we describe a simple, versatile method to enhance differentiation of multiple mesendoderm-derived cells types with a brief pre-differentiation protocol. After 48 hours of treatment with moderate concentrations of both BMP4 and Activin A (referred to as BA), a marked reduction of pluripotency genes and proteins was observed, concurrent with an up-regulation of mesendoderm genes and their protein products. Transcriptomic analysis revealed that by 48 hours, cells induced with BA up-regulated genes associated with a range of mesoderm and endoderm cell lineages. Integration of this 48-hour treatment protocol into existing differentiation protocols enhanced the production of chondrocytes and endothelial progenitor cells while reducing neural differentiation capacity. Prolonged exposure of BA treated cells to basal media without exogenous cues for 14 days resulted in single-cell gene expression profiles consistent with mesoderm and endoderm induction. Teratomas formed from cells pre-treated with BA consisted of a higher ratio of mesoderm and endoderm to ectoderm tissue than teratomas formed from untreated cells. Thus, our pre-differentiation system is a simple and effective means for production of mesendoderm progenitors as well as downstream cell lineages.

## Results

### Single-cell seeding in defined conditions produces robust mesendoderm differentiation

The addition of BMP4 and Activin A to basal media has been shown to induce differentiation of hPSCs to a primitive streak/mesendoderm phenotype in standard colony-seeded cultures (Teo et al., 2012), and more recently, in a micropatterned colony culture system (Nazareth et al., 2013). We sought to approximate the rigorous spatial control afforded by the micropatterned system and the subsequent control over paracrine signaling effects while circumventing the microcontact printing step necessary to produce micropatterns. We hypothesized that the stricter control of initial cell density in single-cell seeding (SCS) would allow for more uniform and reproducible cell dispersions than colony seeding, which would in turn produce more rapid and robust mesendoderm differentiation, similar to micropatterned cultures. To test this hypothesis, we assessed the spatial uniformity of cells seeded by colony and SCS methodologies as well as the downstream differentiation response.

Chemically defined, feeder-free conditions comprising Essential 8 (E8) media (Chen et al., 2011) and Matrigel-coated dishes were used for hPSCs maintenance to reduce lot and batch variability associated with serum and feeder cells, and to optimize hPSC homogeneity. Colonies were seeded by splitting at 1:9, 1:6, and 1:3 ratios, while SCS was done at densities of 1.0, 1.5 and 2.0×105 cells/mL. After overnight seeding, cells were stained with Hoechst (Figure 1A), and high-content imaging was used to assess the uniformity of cell seeding. Coefficient of variation (CV) for the number of cells in a 345×345 μm grid (equivalent to 25 grids in a 10X image) was calculated and normalized to the CV of an equal number of cells in a simulated random uniform distribution to assess uniformity of seeding. SCS resulted in significantly lower normalized CV values than colony seeding across all densities evaluated (p<0.0001), indicating a greater degree of spatial uniformity (Figure 1B). Interestingly, the colony split ratio and SCS density did not have a significant effect on spatial distribution when normalized to a uniform distribution of the same cell number. Thus, SCS produced a spatially uniform cell distribution at a broad range of seeding densities.

**Figure 1.**
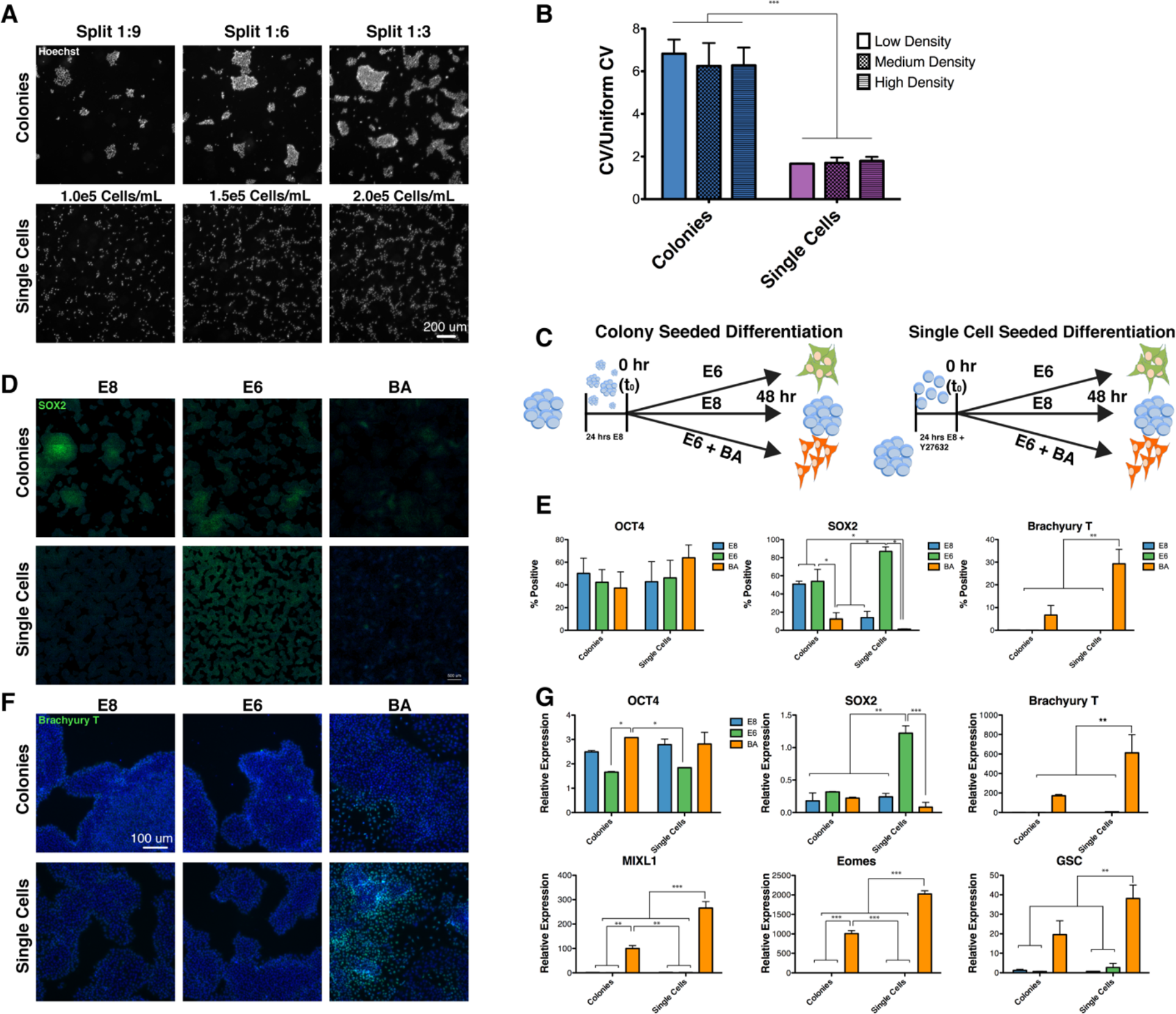
Single cell seeding and BMP4 and Activin A treatment enhance mesendoderm differentiation. (A) hESCs were seeded as colonies at different split ratios or as single cells with different cell densities. Nuclei stained with Hoechst were imaged to quantify the spatial positioning of each cell. Scale bar = 200 μm. (B) The spatial uniformity of cells imaged in (A) was assessed by the coefficient of variation (CV) in the number of cells per grid for a 5×5 pattern overlaid on each field. Error bars represent SEM, n=3, ***=p<0.0001 for colony vs single-cell seeding. (C) Schematic depiction of the colony and single-cell seeded differentiation protocol. (D) Whole-well view of SOX2 staining after 48 hours of differentiation in E8, E6 or BA conditions following colony or single-cell seeding. Scale bar=1 mm. (E) Quantification of immunofluorescent staining for OCT4, SOX2 and T after 48 hours via high-content imaging. Error bars represent SEM, n=2, *=p<0.05, **=p<0.01. (F) Immunofluorescent staining of T after 48 hours of differentiation. Scale bar = 100 μm. (G) Quantification of pluripotency (*OCT4, SOX2*) and mesendoderm (*T, MIXL1, EOMES, GSC*) gene expression by qPCR analysis. Errors bars represent SEM, n=3, *=p<0.05, **=p<0.01, ***=p<0.001.

We then reasoned that the enhanced uniformity of SCS would result in more robust mesendoderm differentiation compared to colony seeding. To test this hypothesis, cells were seeded overnight E8 as colonies (split 1:6) or single cells (1.5×10^5^ cells/mL) and allowed to differentiate spontaneously via removal of TGF-β and FGF2 from E8 media to produce a basal media known as E6, or were directed to mesendoderm by supplementation of BMP4 and Activin A to E6 media (termed BA; schematically depicted in Figure 1C). Cells maintained in E8 served as a pluripotency control, and differentiation was assessed after 48 hours by single cell protein expression and population-based gene expression analysis. OCT4 protein abundance, quantified by immunofluorescence followed by high-content imaging, was not significantly different for any combination of treatment (E8, E6, BA) or seeding method (colony, single cell). The percentage of SOX2-positive cells, which is indicative of both pluripotent and neuroectodermal cells, was significantly higher in single cell E6 cultures compared to all other treatments (Figure 1D, E). Interestingly, the spatial distribution of SOX2 in SCS E6 cultures appeared uniform and homogeneous, whereas SOX2 expression in colony E8 and E6 cultures was notably more heterogeneous, with clusters of positive and negative cells (Figure 1D). Virtually no expression of SOX2 in SCS BA was observed, suggesting a loss of pluripotency as well as a lack of neuroectoderm differentiation. Expression of Brachyury (T), a marker of primitive streak and mesendoderm differentiation, was significantly higher in SCS BA cultures compared to all other conditions (Figure 1 D,E). T expression in E8 and E6 cultures (both colony and SCS) was nearly zero, indicating that spontaneous mesendoderm differentiation was not observed within this timeframe.

Protein expression data was corroborated by population-level qPCR analysis (Figure 1G). *OCT4* expression was largely unchanged among all treatments, with significant differences observed only between E6 (SCS and colony) and colony BA. Expression of *SOX2* transcripts was significantly higher in SCS E6 than all other treatments, in agreement with observed immunofluorescence staining patterns. In addition to *T*, the expression of early mesoderm and endoderm markers including *MIXL1*, *EOMES*, and *GSC* were assessed by qPCR. With the exception of *GSC*, which was not significantly upregulated compared to colony BA, all of these markers showed higher expression levels in SCS BA compared to all other treatments. Collectively, these data demonstrate that SCS of hPSCs improves the uniformity of spatial dispersion and enhances both spontaneous neuroectoderm differentiation (E6) and directed mesendoderm differentiation (BA) compared to colony seeding.

### BA pre-differentiation enhances mesendoderm differentiation

After ascertaining that SCS could be used to produce uniform and robust differentiation of hPSCs, we next sought to assess global transcriptomic changes in cells treated with E8, E6 and BA by RNA sequencing (RNA-seq). A common undifferentiated sample (i.e. t0) as well as 48-hour E8, E6, and BA samples were sequenced, along with a 24-hour BA time point. Biological replicates clustered closely together at each time point for each of the three treatments, as indicated by hierarchical clustering (Figure 2A). Mesendoderm genes were found to be strongly up-regulated in the 24 and 48-hour time points following BA treatment, including *T*, *EOMES*, *GATA5*, and *MIXL1* (Figure 2A, red boxes), while neuroectoderm-associated genes were strongly down-regulated, including *HES3*, *HTR1A*, *EMX1,* and *LRAT* (Figure 2A, blue box). Enriched Gene Ontology (GO) terms for genes up-regulated in E6 samples and BA samples were identified (Figure 2B). Terms associated with ion channel regulation and nervous system development were enriched in the E6 samples, suggesting E6 medium is permissive of a neuro-ectoderm fate specification. In contrast, terms associated with general differentiation (embryo development/morphogenesis, tissue/organ morphogenesis) as well as mesoderm specific differentiation (circulatory/cardiovascular/blood vessel development, heart development) were strongly enriched at both 24 and 48 hours in the BA treated cells. Additionally, gene set enrichment analysis (GSEA) was performed for the 48-hour BA samples (Figure 2C). Gene sets related to mesendoderm differentiation, including Mesendoderm, Lateral Plate Mesoderm, and Pre-Cartilage Condensation were significantly enriched (p<0.03), while the Neural Ectoderm gene set was not enriched (p=.164). Together, GSEA and GO analysis demonstrate that SCS BA treatment induced a gene expression signature indicative of mesendoderm differentiation, while E6 treatment induced early neuroectoderm specification.

**Figure 2.**
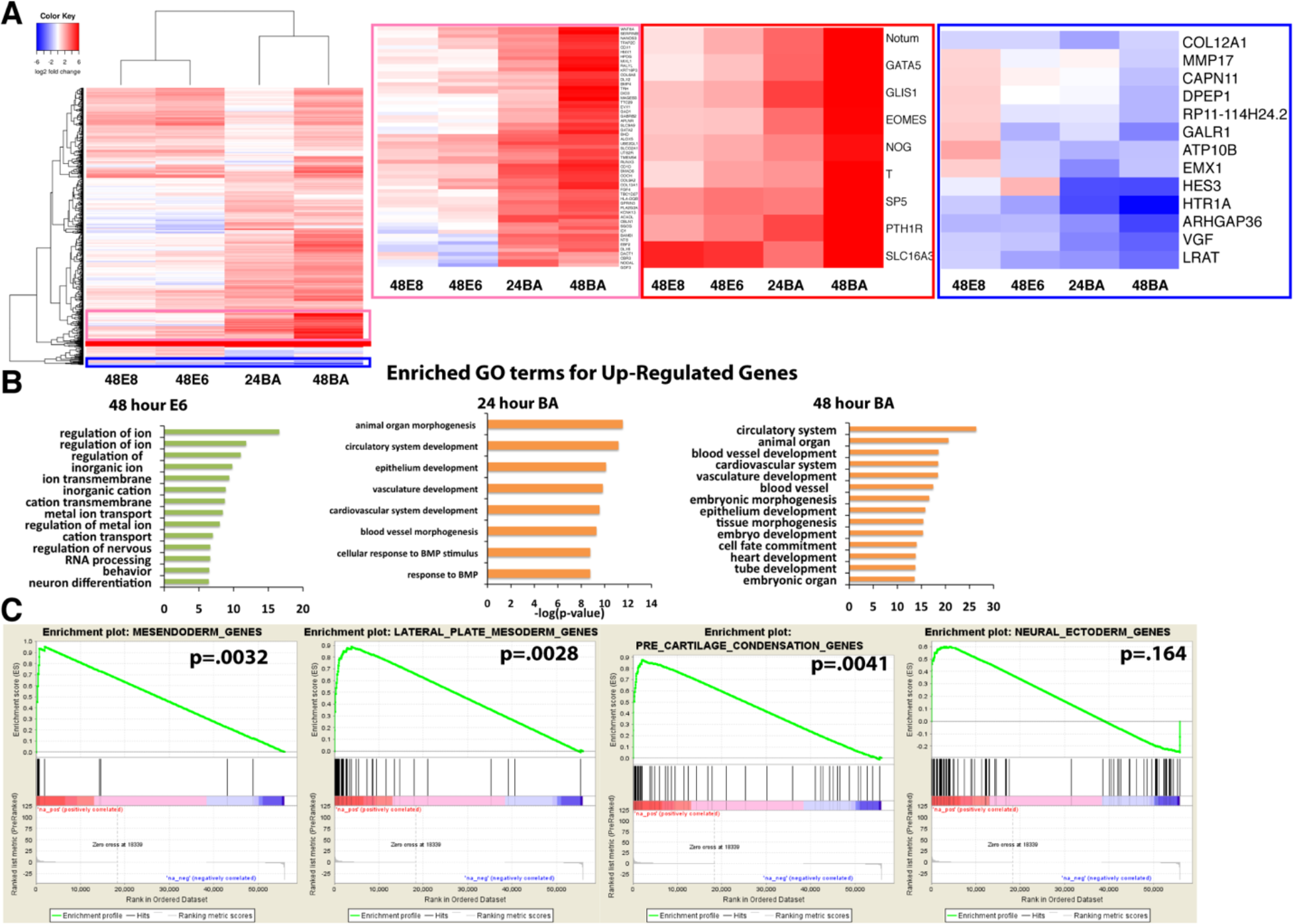
Transcriptomic analysis of E8, E6 and BA treatments by RNA-seq. (A) Heat map of differentially expressed genes between 48-hour E8, 48-hour E6, and 24- and 48-hour BA samples. Heat maps of selected clusters of genes up-regulated in BA (pink box), strongly up-regulated in BA (red box) or down-regulated in BA samples (blue box) are enlarged. (B) Enriched GO terms for genes up-regulated in 48-hour E6, 24-hour BA, and 48-hour BA samples. (C) Gene Set Enrichment Analysis (GSEA) of 48-hour BA samples for Mesendoderm, Lateral Plate Mesoderm, Cartilage Condensation, and Neural Ectoderm gene sets.

### Dynamic transcriptional networks regulate mesendoderm specification

While transcriptomic analysis after 1 and 2 days of differentiation identified distinct gene expression profiles in the three treatment groups, we hypothesized that a higher resolution kinetic analysis would reveal deeper insight into mesendoderm commitment. At 6-hour intervals, samples were collected in the three differentiation conditions for the duration of the 48-hour time course, and RNA-seq was performed. While E8 and E6 samples clustered randomly, the BA samples all clustered sequentially from 6-48 hours, as indicated by hierarchical clustering (Figure 3A, full fold change data in Supplemental Table 4). This observation is further supported by principal component analysis (PCA), with random grouping of E6 and E8 time points observed, contrasting an ordered trajectory of BA samples in the first two PC dimensions (Figure 3B).

**Figure 3.**
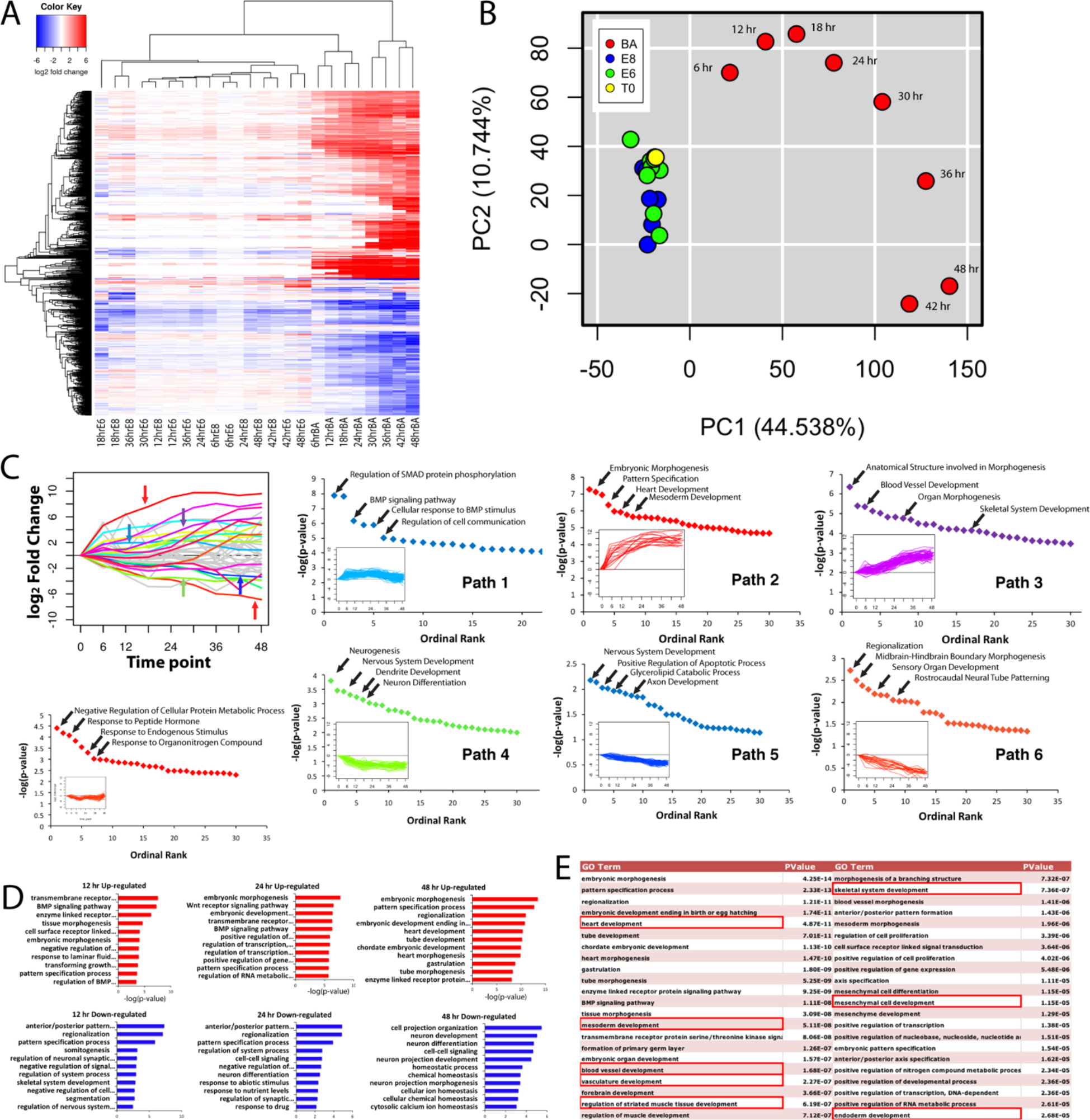
Time course transcriptomic analysis for E8, E6 and BA samples at 6-hour intervals for 48 hours. (A) Heat map of differentially expressed genes, with hierarchical clustering sequentially grouping each BA time point. (B) PCA showing the E8 and E6 samples clustering together, while the BA samples display an ordered trajectory. (C) Genes differentially expressed in BA samples were clustered into paths based on similarity of temporal expression. Genes comprising each path were analyzed for enriched GO terms for upward trajectories (top row), downward trajectories (bottom row), and a trajectory with upward and downward components (bottom left panel). (D) Enriched GO terms for genes up-regulated (top row, red bars) and down-regulated (bottom row, blue bars) in BA samples at 12-, 24- and 48-hour time points. (E) List of enriched GO terms for genes up-regulated at 48 hours in BA samples, with terms related to mesoderm and endoderm differentiation highlighted.

To functionally categorize dynamic gene expression events that regulate mesendoderm differentiation, differentially expressed genes were clustered into discreet paths based on similarity of expression kinetics (Figure 3C). The unique gene sets comprising each path were then queried individually for enriched GO terms. Genes that were upregulated early but quickly plateaued in expression level (Path1) enriched GO terms related to response to growth factor stimulus, such as regulation of SMAD phosphorylation and BMP signaling (full GO analysis can be found in Supplemental Table 1). Similarly, a query of all genes upregulated in BA after 12 hours also enriched terms related to signaling pathways (Figure 3D). Genes that were upregulated at early time points and continued to be strongly upregulated throughout differentiation (Path 2) enriched GO terms related to general differentiation events such as embryonic morphogenesis, but also more specific terms such as heart development. Genes with expression trajectories that increased at later stages of differentiation (Path 3) also enriched terms related to specific differentiation events, such as blood vessel development, skeletal system development, and organ morphogenesis. Collectively, analysis of these upward trajectories indicates that transcriptional response to SCS BA induction occurs in waves, whereby cells initially respond to changes in the signaling environment, followed by general differentiation and morphogenesis, and finally specific differentiation events. Genes clustered in paths with a downward trajectory (Paths 4, 5, 6) enriched terms related to nervous system and neural development and differentiation. This enrichment was similar to that observed with query of genes upregulated at 48 hours in E6 (Figure 2B) as well as genes downregulated at 48 hours in BA (Figure 3D). Broader investigation of all significantly enriched biological process GO terms for genes upregulated at 48 hours following BA treatment revealed many terms related to specific mesoderm and endoderm cell lineages, including heart development, mesoderm development, blood vessel development, regulation of muscle development, mesenchymal cell differentiation, and endoderm development (Figure 3E). These data indicate that after 48 hours of BA treatment, cells may have the potential to be further instructed to differentiate to a variety of mesoderm and endoderm cell types, thereby making the SCS BA protocol amenable to a breadth of tissue engineering applications. However, as the transcriptomic analysis was performed at the population level, the observed gene expression signatures may be the result of either a homogeneous cell population expressing a variety of mesendoderm genes concurrently, or heterogeneous sub-populations each expressing different genes, indicative of divergent differentiation capacities.

### Heterogeneous populations of mesoderm and endoderm cells are produced from initial single cell BA treatment

To address the potential for population heterogeneity contributing to transcriptomic analysis, differentiation at the single-cell level was assessed via single-cell qPCR using a 96-gene panel of pluripotency and differentiation markers on the Fluidigm platform (Figure 4A). Similar to population-level RNA-seq data, cells grown in E8, E6 or BA conditions for 48 hours clustered into distinct groups, as revealed by hierarchical clustering analysis (Figure 4B). E6 and E8 cells clustered close to one another, whereas all BA cells formed a distinct branch separate from the E6 and E8 branches. Similarly, distinct populations of cells between the three treatment groups were observed on a t-distributed stochastic neighbor embedding (t-SNE) plot (Figure 4C), suggesting distinct transcriptional profiles exist for each treatment at the single-cell level. The cluster of genes that distinguished BA cells from E6 and E8 (red box), including *T*, *APLNR*, *GATA6*, *PDGFRA*, *CER1*, *GSC*, and *EOMES*, was enriched for tissues associated with mesoderm lineages (Figure 4D).

**Figure 4.**
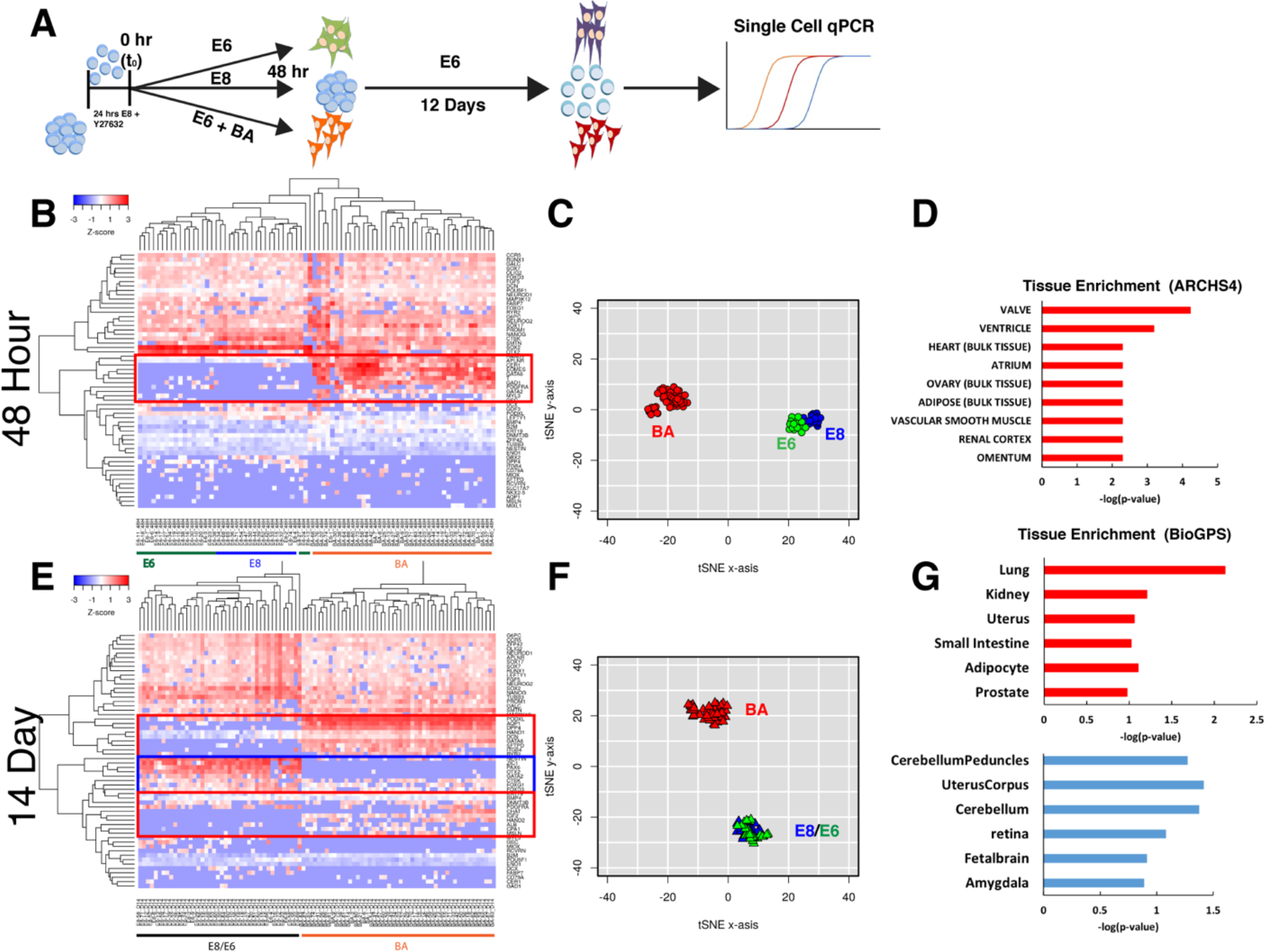
Single cell gene expression analysis. (A) Cells were differentiated for 48 hours in E8, E6, or BA, followed by an additional 12 days in E6 media. Single cell qPCR was performed after 48 hours and 14 days. (B) Heat map of expression levels of the panel of genes in individual cells after 48 hours in E8, E6 or BA. Genes up-regulated in BA cells are highlighted (red box). (C) t-SNE plot of individual cells, with E8 cells in blue, E6 green and BA red. (D) Genes in the red box were analyzed for tissue enrichment using EnrichR. (E) Heat map of expression levels of the panel of genes in individual cells after 14 days. Genes up-regulated in BA cells are highlighted in red boxes and genes up-regulated in E8 and E6 cells are highlighted in the blue box. (F) t-SNE plot of individual after 14 days (E8 blue, E6 green, BA red). (G) Genes up-regulated in BA cells (red boxes in (E)) were analyzed for tissue enrichment in EnrichR (top, red bars), as were genes up-regulated in E8/E6 cells (blue box in (E), bottom graph with blue bars).

To assess the differentiation potential of cells pre-differentiated in E8, E6 or BA, pre-treated cells were allowed to differentiate in basal conditions (E6 or E6 supplemented with B27) for a total of two weeks (48 hours + 12 additional days). The permissive environment afforded by basal conditions allowed cells to differentiate spontaneously along a trajectory dictated by the pre-differentiation conditions alone. Single-cell analysis of differentiation at the 2-week time point was assessed using the same 96-gene panel as the 48-hour time point. Similar to the 48-hour time point, BA-treated cells clustered separately from E6 and E8 cells by both hierarchical clustering (Figure 4E) and t-SNE (Figure 4F). Interestingly, E6 and E8 samples clustered amongst each other, suggesting that pre-differentiation in E6 is not sufficient to alter the long-term differentiation trajectory of cells in basal conditions. A number of genes were highly expressed in the BA-treated population and not expressed in the E6/E8 populations, including *HAND1*, *DCN*, *GATA6*, *AQP1*, *ITGB4*, and *DPP4* (Figure 4E, red boxes), suggestive of mesoderm and endoderm cell lineages. Tissue enrichment analysis for the genes up-regulated in 14-day BA cells identified mesoderm and endoderm derived tissues as most significantly enriched, including lung, kidney, and uterus (Figure 4G, top). Genes that were highly expressed in E6/E8 samples but not BA samples included *PAX6*, *OTX2*, *ZIC1*, *NESTIN*, *GATA2*, *TUBB3*, *SOX2* and *OLIG2* (Figure 4E, blue box), strongly suggestive of a neuroectoderm phenotype. Enriched tissue analysis for these genes up-regulated in E6/E8 cells identified largely brain and ectoderm-related terms (Figure 4G, bottom). The single-cell gene expression observed is consistent with the dense neurite projections and connectivity that was observed in the E6 and E8 samples, but not BA, after 14 days and indicates that pre-differentiation in E6/E8 allows cells to follow a default trajectory to a neuroectoderm fate. Thus, pre-differentiation in BA is permissive to downstream differentiation to both mesoderm and endoderm lineages but diminishes ectoderm potential.

### Pre-differentiation in BA enhances chondrocyte progenitor and endothelial progenitor cell commitment

Population transcriptomic analysis and single-cell qPCR data demonstrated that SCS BA differentiation produced a cell population with a gene expression profile indicative of mesoderm and endoderm derived cells, and previous studies have utilized an adapted version of our protocol to produce skeletal muscle progenitor cells (Shelton et al., 2014, 2016). Therefore, we hypothesized that cells treated for 48 hours in BA could be subsequently specified into mature cell types using existing protocols with enhanced efficiency. The Cartilage Condensation gene set was shown to be significantly enriched after 48-hour BA treatment (Figure 2C), and GO terms related to blood vessel and vasculature development were also found to be significantly enriched following BA treatment (Figure 3C, E). Based on this transcriptomic analysis, existing protocols for chondrogenesis and endothelial progenitor cell (EPC) differentiation were targeted to be modified to include single cell seeding and 48-hour BA pretreatment. For chondrocyte induction, the commonly used micromass culture method was adapted to include the 48-hour pretreatment protocol (Toh et al., 2009) as depicted in Figure 5A. Sulfated glycosaminoglycan (s-GAG) and collagen levels (hydroxy-proline) were quantified after 7 days of micromass culture, and both s-GAG levels and hydroxy-proline expression were significantly higher following BA pre-treatment compared to E8 and E6 controls (Figure 5B). Proteoglycan production, visualized via Alcian Blue staining, was also notably more pronounced in BA-pretreated cells than either E8 or E6 treatments (Figure 5C). The increase in glycosaminoglycan and collagen production was consistently observed for H9 hESCs and two iPSC lines, demonstrating not only that single cell BA pretreatment is amenable to a chondrogenic protocol, but that it significantly enhances differentiation compared to standard differentiation.

**Figure 5.**
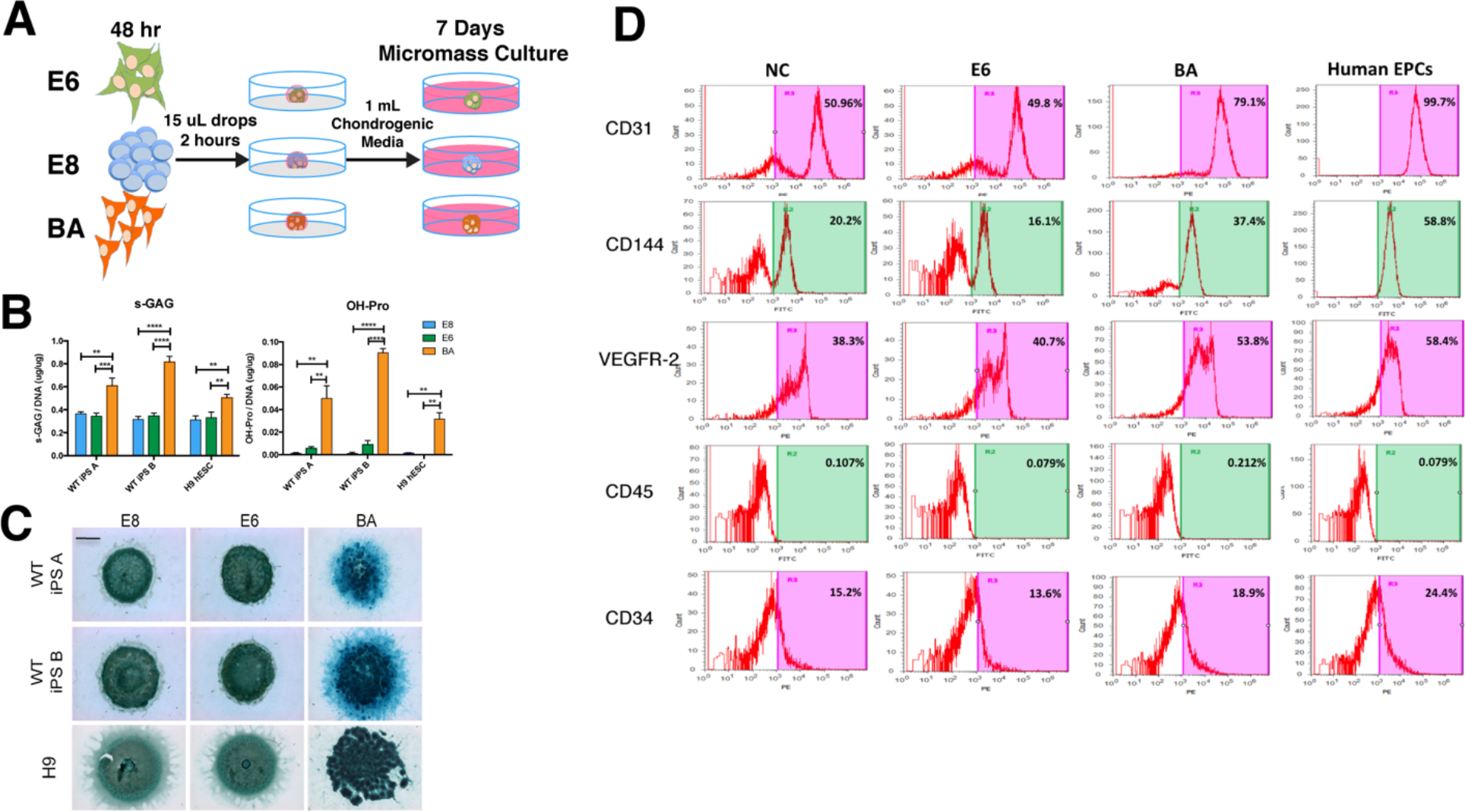
Chondrocyte and endothelial progenitor cell differentiation following BA pretreatment. (A) Schematic depicting the micromass culture protocol utilizing E8, E6 or BA pre-treatment. (B) Sulfated glycosaminoglycan (left) and hydroxy-proline (right) levels were elevated in BA treated cells compared to both E6 and E8 in two iPSC lines and H9 hESCs after 7 days of micromass culture. Errors bars represent SEM, n=3, **=p<0.01, ***=p<0.001, ****=p<0.0001. (C) Alcian blue staining for proteoglycan production after 7 days of micromass culture. Scale bar=2 mm. (D) Flow cytometry analysis of cell surface marker expression after integration of E8 (NC), E6 or BA pretreated cells into an endothelial cell differentiation protocol. Human endothelial progenitor cells are shown as a positive control.

An EPC differentiation protocol was similarly modified to include E8, E6 or BA pretreatment (Tatsumi et al., 2011). To assess EPC differentiation, expression of a number of surface markers was quantified using flow cytometry (Figure 5D). After 6 days of differentiation, BA pretreated cells contained a higher proportion of CD31, CD144, and VEGFR-2 positive cells than E6 pretreated or non-treated controls, indicating higher percentages of endothelial cell differentiation. This again demonstrates that the BA population at 48 hours can be efficiently directed to different cell types given the proper instructive cues.

### BA treatment suppresses spontaneous and directed neuroectoderm differentiation

Enriched GO terms for genes down-regulated upon BA treatment were frequently associated with neural differentiation (Figure 3D), and Neural Ectoderm genes were negative for enrichment in GSEA (Figure 2C). We therefore hypothesized that after pretreatment in BA for 48 hours, cells would be refractory to neural induction. To test this hypothesis, cells treated with E8, E6 or BA were subjected to a common neural differentiation protocol that utilizes dual SMAD inhibition via treatment with LDN193189 and SB431542, inhibitors of BMP and TGF-β, respectively (Chambers et al., 2009). After 48 hours of pre-differentiation (E8, E6 or BA), media was replaced with either E6 alone, or E6 containing LDN and SB and cultured for 3 additional days (Figure 6A). At the 48-hour time point, cells treated with E8 or E6 expressed SOX2 (80% and 88%, respectively), while BA cells showed virtually no expression (2.5% SOX2^+^; Figure 6B, C). Cells treated with E8 expressed high levels of *OCT4*, *NANOG* and *SOX2* transcripts at the 48-hour time point (Figure 6D), confirming that these cells remain in a pluripotent state, whereas the E6 treatment resulted in high expression of *SOX2* and *OTX2* with reduced *OCT4* and *NANOG*, suggesting a loss of pluripotency and early neuroectoderm commitment (Figure 6D). Mesendoderm commitment of BA treated cells was confirmed, as T was expressed exclusively in BA treated cells (Figure 6B), consistent with Figure 1E. Additionally, the loss of *NANOG* and *SOX2* expression along with high expression levels of *OCT4* and *MIXL1* further confirm mesendoderm commitment of BA cells at 48 hours (Figure 6D). Following 5 days of differentiation in both E6 and dual inhibition, pluripotency was lost in all conditions, as a total abrogation of *OCT4* and *NANOG* expression was observed (Figure 6D). High levels of SOX2 expression (both protein and transcript) in E8 and E6 samples was observed, whereas significantly lower SOX2 expression was observed for BA samples in both types of media (Figure 6B,D). Similarly, the intermediate filament protein Nestin was expressed significantly higher in E8 and E6 pretreated cells in both E6 and dual inhibition media. While spontaneous neuroinduction (E6) was nearly absent in BA pretreated cells (1% SOX2^+^, 9% Nestin^+^), higher levels of directed neuroinduction (dual inhibition) was observed (8% SOX2^+^, 32% Nestin^+^), suggesting that a small subset of BA cells remained uncommitted and susceptible to neural differentiation. Interestingly, despite a notable population of Nestin^+^ cells in dual inhibition BA conditions by single cell protein analysis, expression of neural differentiation genes, including *SOX2*, *PAX6*, *OTX2*, and even *Nestin* was extremely low. Finally, BA pretreated cells allowed to spontaneously differentiate in E6 expressed high levels of *AFP* and *KDR*, demonstrating differentiation of mesoderm and endoderm lineages from the 48-hour mesendoderm population.

**Figure 6.**
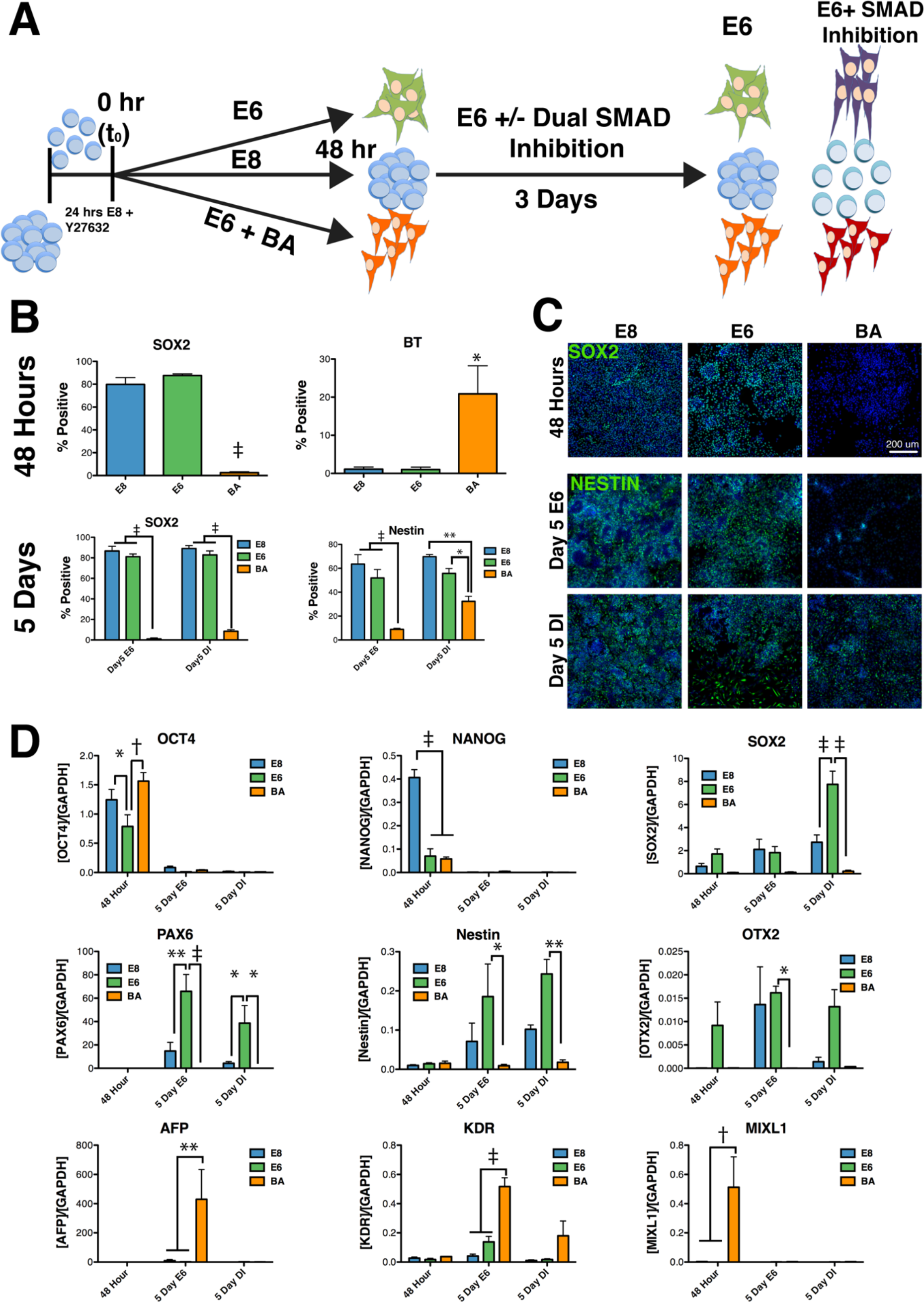
Pretreatment with BA for 48 hours repressed neuroectoderm potential. (A) Schematic depicting the neuro-induction protocol using E8, E6, or BA pre-treated cells is the input population. (B, C) Quantification (B) and representative images (C) of immunofluorescent staining for SOX2 and T after 48 hours, and SOX2 and Nestin after 5 days. (D) Gene expression analysis after 48 hours, 5 days in E6, and 5 days in dual SMAD inhibition media for E8, E6 and BA pretreated cells. Pluripotency (*OCT4*, *NANOG*, *SOX2*), neuroectodoerm *(SOX2*, *PAX6*, *NESTIN*, *OTX2*), and mesoderm/endoderm (*AFP*, *KDR*, *MIXL1*) genes were assessed. Error bars represent SEM, n=3, *=p<0.05, **=p<0.01, ‡=p<0.001. Scale bars=200 μm.

### Teratomas preferentially form mesoderm and endoderm lineages following BA treatment

Cells subjected to SCS BA treatment demonstrated efficient induction of mesendoderm after 48 hours, enhanced efficiency of differentiation of mesoderm lineages, and reduced capacity for neuroectoderm specification under defined conditions *in vitro*. Thus, we hypothesized that in an *in vivo* environment that is permissive to the formation of all three germ lineages (i.e. teratoma formation), BA pretreated cells would preferentially differentiate to mesoderm and endoderm lineages and reduced ectoderm lineages compared to E8 and E6 treatment. To test this hypothesis, cells pretreated with E8, E6 or BA for 48 hours were injected into the hindlimbs of *NOD/SCID* mice to induce teratoma formation. After 9-18 weeks of teratoma growth, tumors were excised, fixed, sectioned, and stained with H&E to identify tissue structures, including cartilage and pigmented epithelium (Figure 7A). The formation of primary germ layers was further assessed via immunofluorescent staining for SOX2 (ectoderm), SOX17 (endoderm) and Desmin (mesoderm) (Figure 7B,C). The ratio of mesoderm and endoderm lineages to ectoderm was calculated for the entire teratoma area of three sections from at least 4 teratomas for each treatment (Figure 7D). Cells pretreated with BA formed teratomas with the highest mesoderm+endoderm to ectoderm ratio, while E6 treatment produced the lowest such ratio (p<0.05 E6 vs. BA). Treatment with E8 resulted in teratomas with a ratio of ∼2 (1.8 ± 0.3), indicating an equal proportion of mesendoderm and ectoderm lineages. Ratios of 4.4 ± 0.8 for BA and 0.8 ± 0.1 for E6 indicate strong bias for mesendoderm and ectoderm differentiation, respectively. Thus, the 48-hour pre-differentiation in BA specifies cells on a differentiation trajectory which, in an *in vivo* environment permissive to formation of germ layers in roughly equal proportions, results in commitment to mesendoderm lineages at the expense of ectoderm formation.

**Figure 7.**
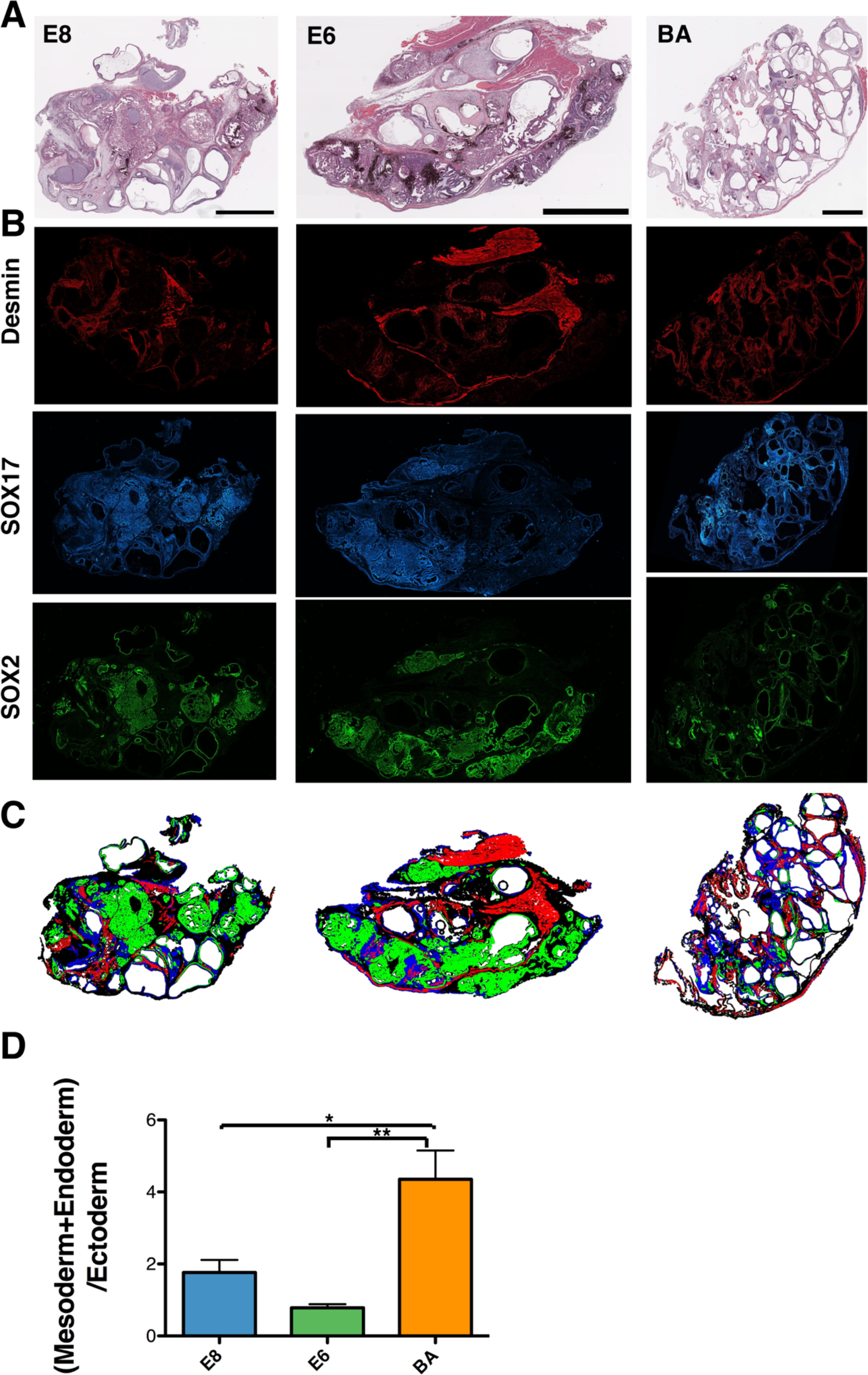
Analysis and quantification of teratomas formed from E8, E6 and BA treated cells. (A) H&E staining of teratomas derived from E8, E6 or BA pre-treated cells. (B) Immunofluorescent staining of teratoma sections for Desmin (mesoderm), SOX17 (endoderm), and SOX2 (ectoderm) was performed and imaged via high content imaging. (C) Images from adjacent sections in (B) were merged to allow quantification of the percentage of each germ layer in each teratoma section. (D) Quantification of the combined mesoderm and endoderm to ectoderm area ratio for teratomas formed from E8, E6 and BA treated cells. For each teratoma, 3 regions at least 150 μm apart were analyzed, and 4 E6, 5 E8, and 6 BA teratomas were used. Scalebars=2 mm. *=p<0.05, **=p<0.005

## Discussion

In this work we describe a simple, robust method for reproducible mesendoderm induction of human PSCs. Transcriptomic data indicated that the cell population produced at 48 hours would be amenable to directed differentiation of cell types including chondrocytes and endothelial progenitor cells, which was confirmed experimentally. We further demonstrated that prolonged exposure to basal media after an initial 48-hour BA induction was sufficient to produce cells with gene expression signatures indicative of mature mesoderm and endoderm cells, whereas E6 induction alone only produced ectodermal gene signatures.

We demonstrate that SCS of PSCs results in quantifiably more homogeneous spatial cellular distribution compared to colony-based seeding. Colony-based seeding strategies can be problematic for reproducibility for a number of reasons. The size and distribution of seeded colonies is a function of the initial colony size, the degree to which colonies are broken up, and the split ratio. Protocols often note the degree of confluency to target before splitting or starting differentiation, which is an approximate and subjective metric dependent on both initial colony-seeding density and colony size. For example, 70% confluence can be achieved by allowing sparsely-seeded colonies to grow very large, or by densely-seeded colonies growing to a smaller final size. Colony-based seeding can also be problematic in communicating the degree to which clumps should be triturated while passaging. The shear stress imparted on clumps of cells during passaging affects the size of clumps seeded, which can in turn affect differentiation trajectory (Bauwens et al., 2008). Thus, pipetting force is an important parameter for a differentiation protocol, yet nearly impossible to objectively articulate in a scientific communication. Furthermore, the issues with confluency and triturating of colonies compound each other, because as described above, 70% confluency can describe colonies of a range of sizes, and therefore even if pipetting shear could be consistent, the starting colony size variance would result in different seeding distributions. By reducing colonies to a single-cell suspension prior to seeding and plating a fixed cell density, we avoided the common issues associated with colony seeding and significantly improved the uniformity and reproducibility of spatial distributions. Furthermore, the parameters we use to define our method are quantitative and objective, allowing others to easily replicate the protocol.

Previous work has demonstrated that numerous combinations of growth factors, small molecules and cytokines can be used to induce mesendoderm differentiation from hPSCs. For example, Touboul et al have used a cocktail consisting of Activin A, FGF2, BMP4, and Ly93092 (termed AFBLy) to induce definitive endoderm (DE) lineages (Touboul et al., 2010), and variations upon this combination to induce other lineages (e.g., FLyA for endoderm, FLyB for mesoderm (Bernardo et al., 2011)). Loh et al have built upon this cocktail and further elucidated the signaling components necessary to induce primitive streak and DE from hPSCs, demonstrating that FGF, BMP and Wnt signaling are required for primitive streak formation (Loh et al., 2014). While the studies mentioned above have all utilized monolayer culture, embryoid body (EB)-based protocols are also commonly used (Craft et al., 2015; Holtzinger et al., 2015; Kattman et al., 2011; Lee et al., 2017; Protze et al., 2016; Witty et al., 2014). EB-based methods for mesoderm induction (cardiac myocytes or chondrocytes) utilize one day of BMP4 treatment followed by an additional two days of BMP4, Activin A and FGF2 treatment (Craft et al., 2015; Kattman et al., 2011; Lee et al., 2017; Protze et al., 2016; Witty et al., 2014). The specific concentration of growth factors is dependent on the targeted cell lineage and the cell line used for induction. Definitive endoderm-derived lineages, including pancreatic cells and hepatocytes, can also be produced from EBs using a similar growth factor cocktail, albeit with higher levels of Activin A (Holtzinger et al., 2015; Nostro et al., 2011).

Our mesendoderm induction protocol shares a number of similarities with these existing methods with notable exceptions. Our findings suggest that endogenous levels of Wnt and FGF signaling – or perhaps pathway crosstalk – are sufficient for mesendoderm induction in the presence of Activin A and BMP4, such that addition of WNT agonists and FGF2 is not required. Interestingly, the specific concentrations of growth factors used varies greatly between protocols. For example, Teo et al reported DE induction with both BMP4 and Activin A concentrations of 50 ng/ml (Teo et al., 2012), whereas many other DE protocols use Activin A levels as high as 100 ng/mL (Bernardo et al., 2011; Brown et al., 2011; Holtzinger et al., 2015; Nostro et al., 2015). For induction of mesoderm, much lower levels of both Activin A and BMP4 have been utilized (1-10 ng/mL, depending on the cell line and target lineage; (Craft et al., 2015; Kattman et al., 2011; Lee et al., 2017; Protze et al., 2016; Witty et al., 2014)). In the presence of FGF2 and PI3K inhibitor (Ly294002), Activin A alone (100 ng/mL) was sufficient to induce DE induction, even in the presence of BMP inhibitor Noggin, while BMP4 alone (10 ng/mL) was able to induce mesoderm in the presence of Activin A inhibitor SB431542 (Bernardo et al., 2011). We demonstrate that 40 ng/mL of both Activin A and BMP4 is ideal for mesendoderm induction in our single-cell seeded culture method. Furthermore, the base media to which growth factors and small molecules are added is quite variable and includes products such as Stem Pro 34 (Craft et al., 2015; Lee et al., 2017; Protze et al., 2016; Witty et al., 2014), RPMI (Nostro et al., 2015; Teo et al., 2012), custom formulations such as chemically defined media (CDM; (Bernardo et al., 2011; Brown et al., 2011; Touboul et al., 2010)), CDM2 (Loh et al., 2014), and serum-free differentiation media (SFD (Holtzinger et al., 2015; Nostro et al., 2011)). Our base differentiation media (E6) has been used to induce neuroectoderm differentiation of hPSCs (Lippmann et al., 2014), in agreement with our results demonstrating that E6 supports neural fate specification. Thus, the context in which signaling molecules are presented to cells appears to contribute to the ultimate response of the cells in terms of cell fate decisions, as various conditions give rise to the same cellular response. Additionally, the conditions under which hPSCs are maintained for self-renewal may contribute to how cells respond to differentiation conditions, as a variety of approaches exist for maintenance, including use of feeder cells (MEFs) and KOSR, MEF-conditioned media, mTESR, E8, as well as a variety of tissue culture coatings including Matrigel and Vitronectin. Transcriptomic studies indicate that variation between pluripotent cell lines is extensive (Adewumi et al., 2007) and that much of the variance can be attributed to the lab in which the cells are cultured (Chin et al., 2009; Newman and Cooper, 2010), suggesting that PSCs are exquisitely sensitive to culture conditions.

We describe a defined differentiation system from which a population of cells with mixed potential emerges. While many studies focus on identifying conditions to direct differentiation to one specific cell type, we identified an intermediate fate from which multiple lineages could be derived. Therefore, our single-cell seeding protocol with BA treatment is a uniquely versatile method for mesendoderm induction that can be integrated into mesoderm and endoderm-derived cell differentiation protocols.

## Experimental Procedures

### Cell Lines and Cell Culture

Human PSCs used in these studies include H9 hESCs and WT iPSCs derived from fibroblasts obtained from the Coriell Institute Biobank (GM00969) (chondrogenic assays) or from late EPCs (Chang et al., 2013) (EPC differentiation assays). Retroviral reprogramming was performed in defined conditions as described previously (Chang et al., 2013). PSCs were maintained in E8 media (Chen et al., 2011) on Matrigel (BD Biosciences)-coated 6-well tissue culture plates, as described previously (Chang et al., 2013). Passaging of hPSCs was done using 0.5 mM EDTA solution as a gentle dissociation agent.

### Colony and Single-cell Seeding and Differentiation

For colony seeding spatial analysis and differentiation, cells were passaged as described above with EDTA and split at 1:3, 1:6, or 1:9 ratios. After overnight colony seeding in E8, media was replaced with fresh E8, E6 or E6 with 40 ng/mL BMP4 and Activin A (R&D Systems or Peprotech). To induce mesendoderm differentiation via single-cell seeding, hPSCs were treated with 10 μm ROCK inhibitor Y-27632 (Tocris) for at least 1 hour prior to dissociating cells with TrypLE Express (Gibco) for 5-7 minutes. An equal volume of E8 containing 15% Knock Out Serum Replacement (KOSR, Gibco) was added and the cell suspension prior to trituration with a P1000. Cells were pelleted by centrifugation (180 rcf, 5 minutes), resuspended in fresh E8 containing 10 μm Y-27632, and a sample was taken for cell counting (Cellometer Auto 2000, Nexcelom Bioscience). Cells were then resuspended at a density of 1.5×105 cells/mL and plated into freshly Matrigel-coated 6 or 12 well tissue culture plates. Seeding densities of 1.0×105-2.0×105 cells/mL were also examined in some experiments. After overnight seeding, E8 media containing Y-27632 was aspirated and replaced with either fresh E8, E6, or E6 with 40 ng/mL BMP4 and Activin A. Cells were allowed to differentiate for 48 hours, with media exchanged after 24 hours.

### Spatial Analysis

Cells were seeded overnight as colonies or single-cells, fixed for 15 minutes in cold 4% paraformaldehyde (PFA), and washed 3x in PBS. Cell nuclei were staining with Hoechst 33342 (Invitrogen) diluted 1:5000 in PBS for 10 minutes. Imaging of fixed and stained cells was performed using the Cellomics ArrayScan VTI (ThermoFisher Scientific) high content imaging instrument. For each condition, the entire well was scanned, the spatial coordinates of each cell in the plate were acquired, and analysis was performed as described in Supplemental Methods.

### RNA-Sequencing

Extraction of RNA was performed using the Macherey-Nagel Nucleospin kit. Concentration and clean-up of RNA was performed via Ethanol/Sodium Acetate precipitation. Library construction and sequencing were performed at the McGill University and Genome Quebec Innovation Centre using the Illumina TruSeq mRNA stranded prep kit, and HiSeq2000 sequencer with 50 or 75bp single end reads. Description of bioinformatics analysis can be found in supplemental material.

### Single Cell Gene Expression

Single cell gene expression analysis was performed using the Fluidigm system. At each time point (48 hours and 14 days), cells were dissociated into a single-cell suspension using TrypLE Express (48 hours) or Dispase (14 days). Isolation, RNA extraction, and cDNA synthesis was performed using the C1 system and C1 Single-Cell Preamp Integrated Fluidic Circuit (IFC) system according to manufacturer’s instructions. Gene expression analysis of amplified cDNA was performed using TAQman gene expression assays (Applied Biosystems) in the BioMark HD on 96.96 Dynamic Array IFCs. Probes are listed in Supplemental Table 2, and description of analysis can be found in Supplemental Methods.

### Mesoderm lineage differentiation and analysis

Complete description of chondrocyte micromass and endothelial progenitor cell differentiation can be found in Supplemental Methods.

### Neural Induction

Single-cell seeding and 48-hour differentiation in E8, E6 or BA was performed as described above. After 48 hours, media was aspirated and replaced with either fresh E6 or E6 supplemented with 10μM SB431542 and 100 nM LDN193189. Media was replaced daily for 3 additional days of differentiation. At the 48-hour and 5-day time points, RNA samples were collected for gene expression analysis, and wells were fixed in 4% PFA for immunofluorescent imaging.

### Quantitative Polymerase Chain Reaction

Gene expression analysis was performed by reverse transcription qPCR using a Roche LightCycler 480. RNA extraction was carried out using the Macherey-Nagel Nucleospin RNA kit and cDNA synthesis was done with Superscript II Reverse Transcriptase (Invitrogen) from 1 μg of RNA. qPCR was performed using cDNA diluted 1:100, 10 μM forward and reverse primers, and 1x Roche LightCycler 480 SYBR Green I MasterMix. Primers are listed in Supplemental Table 3.

### Immunostaining

Cells in 6 or 12 well plates were fixed in cold 4% PFA, as described above. Fixed cells were blocked and permeabilized in 2% Bovine Serum Albumin (BSA) containing 0.01% TritonX-100 for 30 minutes. Primary antibodies were added overnight at 4°C at the dilution or concentration indicated below. After 3 PBS -/-washes, AlexaFluor secondary antibodies diluted 1:200-1:400 were incubated for 1 hour at room temperature. Cell nuclei were stained with Hoechst diluted 1:5000 in PBS for 10 minutes at room temperature. Imaging was performed using the Cellomics ArrayScan. Primary antibodies used are listed in Supplemental Methods.

### Teratomas

Cells pre-differentiated in E8, E6 or BA were dissociated by TrypLE, and 1.0×10^6^ cells were embedded in Matrigel and injected into the tibialis anterior muscles of *NOD/SCID* mice (Charles River Laboratory). These procedures were approved by the University of Ottawa Animal Care Veterinary Services (protocol #OHRIT-1666). Tumors were allowed to form for 9-18 weeks before teratomas were excised, fixed in 4% formaldehyde, and embedded in paraffin. Paraffin-embedded teratomas were sectioned and stained with hematoxylin and eosin (H&E) or by immunofluorescence (IF). For IF, sodium citrate/pressure cooker antigen retrieval was performed. Slides were blocked and permeabilized in 2% BSA, 0.01% Triton X-100 prior to overnight primary antibody incubation at 4° C. Details for SOX2, SOX17 and Desmin antibodies can be found in Supplemental Methods. Secondary antibodies were added for 1 hour at room temperature (AlexaFluor 680 or 488, 1:400). Nuclei were stained with Hoechst 33342 for 15 minutes prior to coverslipping. Imaging of H&E sections was performed using an Aperio CS2 scanscope (Leica Biosystems), and IF sections were imaged using the Cellomics ArrayScan. Quantification of cartilage regions in H&E sections and positive staining in IF was performed using custom scripts written for ImageJ.

### Statistical Analysis

Statistical significance was determined using one-way or two-way ANOVA with Tukey or Bonferroni post-hoc tests using GraphPad Prism software. Three biological replicates were used, except where indicated otherwise.

**RNA-seq data will be submitted to GEO. Accession number will be made available upon request.**

## Author Contributions

RLC designed and performed experiments, analyzed and interpreted data and wrote the manuscript. SYK performed chondrogenic differentiation and analysis, MH and DJS performed EPC differentiation and analysis, CD and JYL assisted with teratoma processing and analysis, CC assisted with single-cell gene expression analysis. RMT and TJP conceptualized and performed bioinformatics analysis. WLS oversaw design and execution of experiments, interpretation of data and wrote the manuscript.

## Acknowledgments

The authors thank the members of the Stanford lab for helpful discussion and review of this project, and Christopher Porter and Gareth Palidwor for bioinformatics assistance. This work is supported by grants from the Canadian Institute of Health Research (MOP-89910), Natural Sciences and Engineering Research Council of Canada (RGPIN 293170-11 & 2016-06081). Infrastructure was supported by the Canadian Foundation for Innovation and the Province of Ontario grants to WLS. WLS is supported by a Canada Research Chair in Integrative Stem Cell Biology.

---

**Supplemental Table 4.** Fold Change analysis of differentially expressed genes for each treatment (E8, E6, BA) and time point (6-48 hour) relative to t_0_ time point.

## References

Adewumi, O., Aflatoonian, B., Ahrlund-Richter, L., Amit, M., Andrews, P.W., Beighton, G., Bello, P.A., Benvenisty, N., Berry, L.S., Bevan, S., et al. (2007). Characterization of human embryonic stem cell lines by the International Stem Cell Initiative. Nat. Biotechnol. 25, 803–816.

Bauwens, C.L., Peerani, R., Niebruegge, S., Woodhouse, K.A., Kumacheva, E., Husain, M., and Zandstra, P.W. (2008). Control of Human Embryonic Stem Cell Colony and Aggregate Size Heterogeneity Influences Differentiation Trajectories. Stem Cells 26, 2300–2310.

Bernardo, A.S., Faial, T., Gardner, L., Niakan, K.K., Ortmann, D., Senner, C.E., Callery, E.M., Trotter, M.W., Hemberger, M., Smith, J.C., et al. (2011). BRACHYURY and CDX2 mediate BMP-induced differentiation of human and mouse pluripotent stem cells into embryonic and extraembryonic lineages. Cell Stem Cell 9, 144–155.

Blin, G., Wisniewski, D., Picart, C., Thery, M., Puceat, M., and Lowell, S. (2018). Geometrical confinement controls the asymmetric patterning of brachyury in cultures of pluripotent cells. Development 145.

Brown, S., Adrian, T.E.O., Pauklin, S., Hannan, N., Cho, C.H.H., Lim, B., Vardy, L., Dunn, N.R., Trotter, M., Pedersen, R., et al. (2011). Activin/nodal signaling controls divergent transcriptional networks in human embryonic stem cells and in endoderm progenitors. Stem Cells 29, 1176–1185.

Chambers, S.M., Fasano, C.A., Papapetrou, E.P., Tomishima, M., Sadelain, M., and Studer, L. (2009). Highly efficient neural conversion of human ES and iPS cells by dual inhibition of SMAD signaling. Nat. Biotechnol. 27, 275–280.

Chang, W.Y., Lavoie, J.R., Kwon, S.Y., Chen, Z., Manias, J.L., Behbahani, J., Ling, V., Kandel, R.A., Stewart, D.J., and Stanford, W.L. (2013). Feeder-independent derivation of induced-pluripotent stem cells from peripheral blood endothelial progenitor cells. Stem Cell Res. 10, 195–202.

Chen, G., Gulbranson, D.R., Hou, Z., Bolin, J.M., Ruotti, V., Probasco, M.D., Smuga-Otto, K., Howden, S.E., Diol, N.R., Propson, N.E., et al. (2011). Chemically defined conditions for human iPSC derivation and culture. Nat. Methods 8, 424–429.

Chin, M.H., Mason, M.J., Xie, W., Volinia, S., Singer, M., Peterson, C., Ambartsumyan, G., Aimiuwu, O., Richter, L., Zhang, J., et al. (2009). Induced Pluripotent Stem Cells and Embryonic Stem Cells Are Distinguished by Gene Expression Signatures. Cell Stem Cell 5, 111–123.

Craft, A.M., Rockel, J.S., Nartiss, Y., Kandel, R.A., Alman, B.A., and Keller, G.M. (2015). Generation of articular chondrocytes from human pluripotent stem cells. Nat. Biotechnol. 33, 638–645.

Davey, R.E., and Zandstra, P.W. (2006). Spatial Organization of Embryonic Stem Cell Responsiveness to Autocrine Gp130 Ligands Reveals an Autoregulatory Stem Cell Niche. Stem Cells 24, 2538–2548.

Dimos, J.T., Rodolfa, K.T., Niakan, K.K., Weisenthal, L.M., Mitsumoto, H., Chung, W., Croft, G.F., Saphier, G., Leibel, R., Goland, R., et al. (2008). Induced Pluripotent Stem Cells Generated from Patients with ALS Can Be Differentiated into Motor Neurons. Science (80-.). 321, 1218–1221.

Draper, J.S., Smith, K., Gokhale, P., Moore, H.D., Maltby, E., Johnson, J., Meisner, L., Zwaka, T.P., Thomson, J.A., and Andrews, P.W. (2004). Recurrent gain of chromosomes 17q and 12 in cultured human embryonic stem cells. Nat. Biotechnol. 22, 53–54.

Evans, M.J., and Kaufman, M.H. (1981). Establishment in culture of pluripotential cells from mouse embryos. Nature 292, 154–156.

Ginis, I., Luo, Y., Miura, T., Thies, S., Brandenberger, R., Gerecht-Nir, S., Amit, M., Hoke, A., Carpenter, M.K., Itskovitz-Eldor, J., et al. (2004). Differences between human and mouse embryonic stem cells. Dev. Biol. 269, 360–380.

Holtzinger, A., Streeter, P.R., Sarangi, F., Hillborn, S., Niapour, M., Ogawa, S., and Keller, G. (2015). New markers for tracking endoderm induction and hepatocyte differentiation from human pluripotent stem cells. Development 142, 4253–4265.

Kattman, S.J., Witty, A.D., Gagliardi, M., Dubois, N.C., Niapour, M., Hotta, A., Ellis, J., and Keller, G. (2011). Stage-Specific Optimization of Activin/Nodal and BMP Signaling Promotes Cardiac Differentiation of Mouse and Human Pluripotent Stem Cell Lines. Cell Stem Cell 8, 228–240.

Lee, J.H., Protze, S.I., Laksman, Z., Backx, P.H., and Keller, G.M. (2017). Human Pluripotent Stem Cell-Derived Atrial and Ventricular Cardiomyocytes Develop from Distinct Mesoderm Populations. Cell Stem Cell 21, 179–194.e4.

Lee, L.H., Peerani, R., Ungrin, M., Joshi, C., Kumacheva, E., and Zandstra, P. (2009). Micropatterning of human embryonic stem cells dissects the mesoderm and endoderm lineages. Stem Cell Res. 2, 155–162.

Lippmann, E.S., Estevez-Silva, M.C., and Ashton, R.S. (2014). Defined Human Pluripotent Stem Cell Culture Enables Highly Efficient Neuroepithelium Derivation Without Small Molecule Inhibitors. Stem Cells 32, 1032–1042.

Loh, K.M., Ang, L.T., Zhang, J., Kumar, V., Ang, J., Auyeong, J.Q., Lee, K.L., Choo, S.H., Lim, C.Y.Y., Nichane, M., et al. (2014). Efficient endoderm induction from human pluripotent stem cells by logically directing signals controlling lineage bifurcations. Cell Stem Cell 14, 237–252.

Martin, G.R. (1981). Isolation of a pluripotent cell line from early mouse embryos cultured in medium conditioned by teratocarcinoma stem cells (embryonic stem cells/inner cell masses/differentiation in vsitro/embryonal carcinoma cells/growth factors).

Mitalipova, M.M., Rao, R.R., Hoyer, D.M., Johnson, J.A., Meisner, L.F., Jones, K.L., Dalton, S., and Stice, S.L. (2005). Preserving the genetic integrity of human embryonic stem cells. Nat. Biotechnol. 23, 19–20.

Mohr, J.C., de Pablo, J.J., and Palecek, S.P. (2006). 3-D microwell culture of human embryonic stem cells. Biomaterials 27, 6032–6042.

Nazareth, E.J.P., Ostblom, J.E.E., Lücker, P.B., Shukla, S., Alvarez, M.M., Oh, S.K.W., Yin, T., and Zandstra, P.W. (2013). High-throughput fingerprinting of human pluripotent stem cell fate responses and lineage bias. Nat. Methods 10, 1225–1231.

Newman, A.M., and Cooper, J.B. (2010). Lab-Specific Gene Expression Signatures in Pluripotent Stem Cells. Cell Stem Cell 7, 258–262.

Niakan, K.K., and Eggan, K. (2013). Analysis of human embryos from zygote to blastocyst reveals distinct gene expression patterns relative to the mouse. Dev. Biol. 375, 54–64.

Nostro, M.C., Sarangi, F., Ogawa, S., Holtzinger, A., Corneo, B., Li, X., Micallef, S.J., Park, I.-H., Basford, C., Wheeler, M.B., et al. (2011). Stage-specific signaling through TGF family members and WNT regulates patterning and pancreatic specification of human pluripotent stem cells. Development 138, 861–871.

Nostro, M.C., Sarangi, F., Yang, C., Holland, A., Elefanty, A.G., Stanley, E.G., Greiner, D.L., and Keller, G. (2015). Efficient Generation of NKX6-1+ Pancreatic Progenitors from Multiple Human Pluripotent Stem Cell Lines. Stem Cell Reports 4, 591–604.

Park, I.-H., Arora, N., Huo, H., Maherali, N., Ahfeldt, T., Shimamura, A., Lensch, M.W., Cowan, C., Hochedlinger, K., and Daley, G.Q. (2008). Disease-Specific Induced Pluripotent Stem Cells. Cell 134, 877–886.

Peerani, R., Rao, B.M., Bauwens, C., Yin, T., Wood, G.A., Nagy, A., Kumacheva, E., and Zandstra, P.W. (2007). Niche-mediated control of human embryonic stem cell self-renewal and differentiation. EMBO J. 26, 4744–4755.

Protze, S.I., Liu, J., Nussinovitch, U., Ohana, L., Backx, P.H., Gepstein, L., and Keller, G.M. (2016). Sinoatrial node cardiomyocytes derived from human pluripotent cells function as a biological pacemaker. Nat. Biotechnol. 35, 56–68.

Reubinoff, B.E., Pera, M.F., Fong, C.-Y., Trounson, A., and Bongso, A. (2000). Embryonic stem cell lines from human blastocysts: somatic differentiation in vitro. Nat. Biotechnol. 18, 399–404.

Shelton, M., Metz, J., Liu, J., Carpenedo, R.L., Demers, S.-P., Stanford, W.L., and Skerjanc, I.S. (2014). Derivation and expansion of PAX7-positive muscle progenitors from human and mouse embryonic stem cells. Stem Cell Reports 3.

Shelton, M., Kocharyan, A., Liu, J., Skerjanc, I.S., and Stanford, W.L. (2016). Robust generation and expansion of skeletal muscle progenitors and myocytes from human pluripotent stem cells.

Sheng, G. (2015). Epiblast morphogenesis before gastrulation. Dev. Biol. 401, 17–24.

Soldner, F., Hockemeyer, D., Beard, C., Gao, Q., Bell, G.W., Cook, E.G., Hargus, G., Blak, A., Cooper, O., Mitalipova, M., et al. (2009). Parkinson’s Disease Patient-Derived Induced Pluripotent Stem Cells Free of Viral Reprogramming Factors. Cell 136, 964–977.

Solnica-Krezel, L. (2005). Conserved Patterns of Cell Movements during Vertebrate Gastrulation. Curr. Biol. 15, R213–R228.

Tabar, V., and Studer, L. (2014). Pluripotent stem cells in regenerative medicine: challenges and recent progress. Nat. Rev. Genet. 15, 82–92.

Takahashi, K., Tanabe, K., Ohnuki, M., Narita, M., Ichisaka, T., Tomoda, K., and Yamanaka, S. (2007). Induction of Pluripotent Stem Cells from Adult Human Fibroblasts by Defined Factors. Cell 131, 861–872.

Tam, P.P.L., and Loebel, D.A.F. (2007). Gene function in mouse embryogenesis: Get set for gastrulation. Nat. Rev. Genet. 8, 368–381.

Tam, P.P., Loebel, D.A., and Tanaka, S.S. (2006). Building the mouse gastrula: signals, asymmetry and lineages. Curr. Opin. Genet. Dev. 16, 419–425.

Tatsumi, R., Suzuki, Y., Sumi, T., Sone, M., Suemori, H., and Nakatsuji, N. (2011). Simple and highly efficient method for production of endothelial cells from human embryonic stem cells. Cell Transplant. 20, 1423–1430.

Teo, A.K.K., Ali, Y., Wong, K.Y., Chipperfield, H., Sadasivam, A., Poobalan, Y., Tan, E.K., Wang, S.T., Abraham, S., Tsuneyoshi, N., et al. (2012). Activin and BMP4 synergistically promote formation of definitive endoderm in human embryonic stem cells. Stem Cells 30, 631– 642.

Thomson, J.A., Itskovitz-Eldor, J., Shapiro, S.S., Waknitz, M.A., Swiergiel, J.J., Marshall, V.S., and Jones, J.M. (1998). Embryonic stem cell lines derived from human blastocysts. Science 282, 1145–1147.

Toh, W.S., Guo, X.M., Choo, A.B., Lu, K., Lee, E.H., and Cao, T. (2009). Differentiation and enrichment of expandable chondrogenic cells from human embryonic stem cells in vitro. J. Cell. Mol. Med. 13, 3570–3590.

Touboul, T., Hannan, N.R.F., Corbineau, S., Martinez, A., Martinet, C., Branchereau, S., Mainot, S., Strick-Marchand, H., Pedersen, R., Di Santo, J., et al. (2010). Generation of functional hepatocytes from human embryonic stem cells under chemically defined conditions that recapitulate liver development. Hepatology 51, 1754–1765.

Wei, C.L., Miura, T., Robson, P., Lim, S.-K., Xu, X.-Q., Lee, M.Y.-C., Gupta, S., Stanton, L., Luo, Y., Schmitt, J., et al. (2005). Transcriptome Profiling of Human and Murine ESCs Identifies Divergent Paths Required to Maintain the Stem Cell State. Stem Cells 23, 166–185.

Witty, A.D., Mihic, A., Tam, R.Y., Fisher, S.A., Mikryukov, A., Shoichet, M.S., Li, R.-K., Kattman, S.J., and Keller, G. (2014). Generation of the epicardial lineage from human pluripotent stem cells. Nat. Biotechnol. 32, 1026–1035.

Wu, S.M., and Hochedlinger, K. (2011). Harnessing the potential of induced pluripotent stem cells for regenerative medicine. Nat. Cell Biol. 13, 497–505.

Yu, J., Vodyanik, M.A., Smuga-Otto, K., Antosiewicz-Bourget, J., Frane, J.L., Tian, S., Nie, J., Jonsdottir, G.A., Ruotti, V., Stewart, R., et al. (2007). Induced Pluripotent Stem Cell Lines Derived from Human Somatic Cells. Science (80-.). 318, 1917–1920.

