## Supplemental Material file 1 for "Transcriptomically-guided mesendoderm induction of human pluripotent stem cells using a systematically defined culture scheme"

### Supplemental Table 1. Complete gene lists and GO terms from Figure 3C.

**Path 1 Genes**: RP11-34P13.15, RP4-758J18.10, VWA1, CHD5, AZIN2, FOXO6, RP11-403I13.8, ARHGAP30, RGS4, LRRN2, RASSF5, SERTAD4, GJC2, RHOU, REEP1, FOXI3, SH3RF3, COL4A4, ZDHHC23, FGFR3, PPP2R2C, CTD-2031P19.4, RNF182, GRM4, PRR15, DGKI, CHMP4C, CALB1, SPAG1, KLF4, ENG, RET, GDF10, ADAMTS14, SPOCK2, MBL1P, ADAM8, LRP4-AS1, CARNS1, DGAT2, CRYAB, AP000783.1, OPCML, PLEKHG6, GDF3, EMP1, RASSF9, FAM101A, STON2, GREM1, ACTC1, CORO2B, FURIN, WFIKKN1, BAIAP3, TMC5, HS3ST4, ZFHX3, NLRP1, RASD1, CACNG4, EMILIN2, L3MBTL4, KLHL14, HMSD, RP11-849I19.1, SALL3, GADD45B, KANK3, CTC-526N19.1, ZNF888, MMP9, BMP7, PIK3IP1, MCHR1, SYTL5, CAMK2N1, PINK1, ID3, PTPRU, MANEAL, MCOLN3, LRRC8C, NTNG1, KCNC4, RP11, 430C7.5, C1orf95, ID2-AS1, ID2, GDF7, KCNG3, RGPD8, PSD4, CCDC74B, BMPR2, KAT2B, LINC00693, ZNF654, FILIP1L, SH3TC1, CPEB2, NPFFR2, TRPC3, RP11-752L20.3, FAM198B, TLL1, CDH9, PDZD2, CHSY3, GALNT10, FOXQ1, ATXN1, ID4, COL11A2, CNR1, GTF2IP4, FZD1, PAX5, RP11-35N6.1, UNC5B, NKX1-2, FAM196A, EBF3, PRRG4, LRP4, SYT7, PLBD1, GRASP, ALX1, HIP1R, LPAR6, SLITRK6, C16orf89, RP11-491F9.1, MMP2, B3GNT9, NXPH3, TNRC6C-AS1, LDLRAD4, NOL4, SMAD7, HCN2, PDE4A, KANK2, SAMD1, EXOC3L2, IL11, EMILIN3, KCNB1, DOK5, EEF1A2, A4GALT, ADGRG2, ELF4, ABCD1

| Term | Count | % | PValue | Genes |
|---|---|---|---|---|
| regulation of pathway-restricted SMAD protein phosphorylation | 9 | 6.34 | 1.31E-08 | GDF3, SMAD7, GDF7, BMPR2, GDF10, GREM1, BMP7, LDLRAD4, ENG |
| pathway-restricted SMAD protein phosphorylation | 9 | 6.34 | 1.50E-08 | GDF3, SMAD7, GDF7, BMPR2, GDF10, GREM1, BMP7, LDLRAD4, ENG |
| BMP signaling pathway | 10 | 7.04 | 6.88E-07 | GDF3, SMAD7, GDF7, BMPR2, FZD1, GDF10, GREM1, BMP7, ENG, LRP4 |
| response to BMP | 10 | 7.04 | 1.24E-06 | GDF3, SMAD7, GDF7, BMPR2, FZD1, GDF10, GREM1, BMP7, ENG, LRP4 |
| cellular response to BMP stimulus | 10 | 7.04 | 1.24E-06 | GDF3, SMAD7, GDF7, BMPR2, FZD1, GDF10, GREM1, BMP7, ENG, LRP4 |
| regulation of cell communication | 43 | 30.28 | 9.74E-06 | GDF3, KCNC4, FGFR3, HIP1R, GDF7, MMP9, BMPR2, PINK1, SYT7, GREM1, RHOU, CALB1, KANK2, IL11, UNC5B, PDE4A, PLEKHG6, CNR1, NPFFR2, ADAM8, CHD5, RET, SMAD7, KCNB1, FZD1, PSD4, CACNG4, DGKI, PIK3IP1, LDLRAD4, FURIN, ARHGAP30, SALL3, GRM4, DOK5, LPAR6, RGS4, GDF10, GADD45B, BMP7, ENG, KLF4, LRP4 |
| ossification | 13 | 9.15 | 1.19E-05 | FGFR3, MMP9, BMPR2, FZD1, GREM1, MMP2, ID2, GDF10, ID4, ID3, BMP7, COL11A2, LRP4 |
| regulation of signaling | 43 | 30.28 | 1.48E-05 | GDF3, KCNC4, FGFR3, HIP1R, GDF7, MMP9, BMPR2, PINK1, SYT7, GREM1, RHOU, CALB1, KANK2, IL11, UNC5B, PDE4A, PLEKHG6, CNR1, NPFFR2, ADAM8, CHD5, RET, SMAD7, KCNB1, FZD1, PSD4, CACNG4, DGKI, PIK3IP1, LDLRAD4, FURIN, ARHGAP30, SALL3, GRM4, DOK5, LPAR6, RGS4, GDF10, GADD45B, BMP7, ENG, KLF4, LRP4 |
| embryonic morphogenesis | 16 | 11.27 | 1.68E-05 | GDF3, RET, GDF7, MMP9, BMPR2, FZD1, PAX5, GREM1, MMP2, ID2, BMP7, SLITRK6, ENG, LRP4, KLF4, ALX1 |
| regulation of phosphorylation | 26 | 18.31 | 1.90E-05 | GDF3, FGFR3, GDF7, MMP9, BMPR2, PINK1, GREM1, IL11, NPFFR2, ADAM8, RET, KAT2B, SMAD7, EEF1A2, FZD1, PIK3IP1, LDLRAD4, CAMK2N1, GRM4, RGS4, GDF10, GADD45B, BMP7, ENG, LRP4, KLF4 |
| transmembrane receptor protein serine/threonine kinase signaling pathway | 12 | 8.45 | 2.30E-05 | GDF3, SMAD7, GDF7, BMPR2, FZD1, GDF10, GREM1, BMP7, LDLRAD4, FURIN, ENG, LRP4 |
| regulation of transmembrane receptor protein serine/threonine kinase signaling pathway | 10 | 7.04 | 2.45E-05 | GDF3, SMAD7, GDF7, BMPR2, FZD1, GDF10, GREM1, BMP7, LDLRAD4, ENG |
| positive regulation of pathway-restricted SMAD protein phosphorylation | 6 | 4.23 | 2.49E-05 | GDF3, GDF7, BMPR2, GDF10, BMP7, ENG |
| extracellular matrix organization | 12 | 8.45 | 3.21E-05 | COL4A4, ADAMTS14, SPOCK2, MMP9, COL11A2, GREM1, VWA1, ADAM8, FURIN, ENG, MMP2, TLL1 |
| extracellular structure organization | 12 | 8.45 | 3.30E-05 | COL4A4, ADAMTS14, SPOCK2, MMP9, COL11A2, GREM1, VWA1, ADAM8, FURIN, ENG, MMP2, TLL1 |

**Path 2 Genes:** RP11-54O7.3, SAMD11, GLIS1, MIXL1, GREM2, SP5, EOMES, PTH1R, HAND1, MSX2, T, EVX1, BHLHE22, TBX3, NOTUM, GATA6, PCAT14

| Term | Count | % | PValue | Genes |
|---|---|---|---|---|
| regulation of transcription from RNA polymerase II promoter | 11 | 68.75 | 5.40E-08 | MSX2, T, BHLHE22, HAND1, TBX3, EVX1, GATA6, GLIS1, EOMES, GREM2, MIXL1 |
| embryonic morphogenesis | 8 | 50 | 7.71E-08 | MSX2, T, HAND1, TBX3, GATA6, EOMES, GREM2, MIXL1 |
| embryo development | 9 | 56.25 | 1.12E-07 | MSX2, T, HAND1, TBX3, EVX1, GATA6, EOMES, GREM2, MIXL1 |
| pattern specification process | 7 | 43.75 | 4.57E-07 | MSX2, T, HAND1, TBX3, EVX1, EOMES, GREM2 |

| | | | | |
|---|---|---|---|---|
| transcription from RNA polymerase II promoter | 10 | 62.5 | 1.04E-06 | MSX2, T, HAND1, TBX3, EVX1, GATA6, GLIS1, EOMES, GREM2, MIXL1 |
| heart development | 7 | 43.75 | 1.16E-06 | MSX2, T, HAND1, TBX3, GATA6, EOMES, MIXL1 |
| formation of primary germ layer | 5 | 31.25 | 1.63E-06 | T, HAND1, GATA6, EOMES, MIXL1 |
| mesoderm development | 5 | 31.25 | 2.19E-06 | T, HAND1, TBX3, EOMES, MIXL1 |
| regulation of transcription, DNA-templated | 12 | 75 | 2.42E-06 | MSX2, T, BHLHE22, HAND1, TBX3, EVX1, GATA6, GLIS1, EOMES, SP5, GREM2, MIXL1 |
| transcription, DNA-templated | 12 | 75 | 2.43E-06 | MSX2, T, BHLHE22, HAND1, TBX3, EVX1, GATA6, GLIS1, EOMES, SP5, GREM2, MIXL1 |
| regulation of nucleic acid-templated transcription | 12 | 75 | 2.58E-06 | MSX2, T, BHLHE22, HAND1, TBX3, EVX1, GATA6, GLIS1, EOMES, SP5, GREM2, MIXL1 |
| regulation of RNA biosynthetic process | 12 | 75 | 2.72E-06 | MSX2, T, BHLHE22, HAND1, TBX3, EVX1, GATA6, GLIS1, EOMES, SP5, GREM2, MIXL1 |
| regionalization | 6 | 37.5 | 3.30E-06 | MSX2, T, TBX3, EVX1, EOMES, GREM2 |
| regulation of RNA metabolic process | 12 | 75 | 3.81E-06 | MSX2, T, BHLHE22, HAND1, TBX3, EVX1, GATA6, GLIS1, EOMES, SP5, GREM2, MIXL1 |
| nucleic acid-templated transcription | 12 | 75 | 3.90E-06 | MSX2, T, BHLHE22, HAND1, TBX3, EVX1, GATA6, GLIS1, EOMES, SP5, GREM2, MIXL1 |

**Path 3 Genes:** CSF3R, PTGER3, IFI16, PRRX1, PLXNA2, DUSP10, AMER3, LINC01124, ADAMTS9, TRH, COL6A6, PLSCR4, JAKMIP1, ARHGAP24, LEF1, CTD-2035E11.4, ANXA2R, AC025171.1, ANKRD55, HAPLN1, GABRB2, FOXC1, SERPINB9, HLA-DQA1, HLA-DQB1, BMPER, TBX20, RELN, KEL, DLC1, DKK4, PSKH2, HAS2, CCDC3, ST8SIA6, COL13A1, FGF8, PPAPDC1A, WNT5B, ATP12A, PCDH17, GSC, SYNE3, CA12, MESP1, ANPEP, NKD1, KRT16P2, MFAP4, AC004448.5, KRT16P3, RP11-445F12.1, TBX4, FADS6, CYGB, SLC16A3, APOBEC3B-AS1, APOBEC3G, NFAM1, MAGEB3, SLITRK2

| Term | Count | % | PValue | Genes |
|---|---|---|---|---|
| anatomical structure formation involved in morphogenesis | 16 | 28.57 | 4.49E-07 | DLC1, NKD1, FGF8, GSC, PLXNA2, TBX20, TBX4, LEF1, ANPEP, ARHGAP24, DKK4, BMPER, CSF3R, RELN, FOXC1, MESP1 |
| regionalization | 9 | 16.07 | 4.05E-06 | NKD1, FGF8, GSC, PLXNA2, TBX20, LEF1, RELN, FOXC1, MESP1 |
| blood vessel development | 11 | 19.64 | 4.54E-06 | FGF8, BMPER, TBX20, TBX4, PRRX1, LEF1, HAS2, FOXC1, ANPEP, ARHGAP24, MESP1 |
| vasculature development | 11 | 19.64 | 7.52E-06 | FGF8, BMPER, TBX20, TBX4, PRRX1, LEF1, HAS2, FOXC1, ANPEP, ARHGAP24, MESP1 |
| blood vessel morphogenesis | 10 | 17.86 | 9.44E-06 | FGF8, BMPER, TBX20, TBX4, PRRX1, LEF1, HAS2, FOXC1, ANPEP, ARHGAP24 |
| localization of cell | 15 | 26.79 | 1.46E-05 | DLC1, NKD1, FGF8, WNT5B, PLXNA2, TBX20, LEF1, SLC16A3, BMPER, CSF3R, RELN, CYGB, FOXC1, HAS2, MESP1 |
| cell motility | 15 | 26.79 | 1.46E-05 | DLC1, NKD1, FGF8, WNT5B, PLXNA2, TBX20, LEF1, SLC16A3, BMPER, CSF3R, RELN, CYGB, FOXC1, HAS2, MESP1 |
| organ morphogenesis | 13 | 23.21 | 1.77E-05 | DLC1, FGF8, NKD1, GSC, COL13A1, TBX20, TBX4, PRRX1, LEF1, CSF3R, FOXC1, HAS2, MESP1 |
| cell migration | 14 | 25.00 | 2.05E-05 | DLC1, FGF8, WNT5B, PLXNA2, TBX20, LEF1, SLC16A3, BMPER, CSF3R, RELN, FOXC1, HAS2, CYGB, MESP1 |
| pattern specification process | 9 | 16.07 | 3.12E-05 | NKD1, FGF8, GSC, PLXNA2, TBX20, LEF1, RELN, FOXC1, MESP1 |
| epithelium development | 13 | 23.21 | 3.42E-05 | DLC1, NKD1, FGF8, WNT5B, GSC, PLXNA2, TBX20, TBX4, LEF1, DKK4, BMPER, FOXC1, MESP1 |
| tube development | 10 | 17.86 | 3.53E-05 | DLC1, FGF8, BMPER, GSC, PLXNA2, TBX20, TBX4, LEF1, FOXC1, MESP1 |
| mesenchyme development | 7 | 12.50 | 5.41E-05 | FGF8, GSC, TBX20, LEF1, HAS2, FOXC1, MESP1 |
| circulatory system development | 12 | 21.43 | 6.04E-05 | DLC1, FGF8, BMPER, TBX20, TBX4, PRRX1, LEF1, HAS2, FOXC1, ANPEP, ARHGAP24, MESP1 |
| cardiovascular system development | 12 | 21.43 | 6.04E-05 | DLC1, FGF8, BMPER, TBX20, TBX4, PRRX1, LEF1, HAS2, FOXC1, ANPEP, ARHGAP24, MESP1 |

**Path 4 Genes:** FAM46B, ACTA1, NXPH2, LRP2, CCDC141, MAP2, UNC80, IGFBP5, STAC, EPHA3, FOXL2NB, TM4SF18, P2RY1, PTX3, SLITRK3, LIMCH1, CXCL5, LRAT, IRX2, GCNT4, FAM65B, C6orf141, HECW1, IGFBP3, VWC2, COBL, SFRP1, NPM1P21, PRDM14, MAMDC2, EIF2S2P3, ANO1, MAP6, ADAMTS8, A2M, PAPLN, GABRA5, GPR176, GRIN2A, SEZ6, CH17-431G21.1, PRKCA, DCC, SLC24A3, KIAA1644, APLN

| Term | Count | % | PValue | Genes |
|---|---|---|---|---|
| neurogenesis | 12 | 26.67 | 1.60E-04 | DCC, COBL, SLITRK3, SFRP1, P2RY1, MAP2, GABRA5, GRIN2A, VWC2, MAP6, SEZ6, EPHA3 |
| type B pancreatic cell proliferation | 3 | 6.67 | 3.49E-04 | SFRP1, IGFBP3, IGFBP5 |
| nervous system development | 14 | 31.11 | 3.76E-04 | DCC, COBL, GABRA5, GRIN2A, EPHA3, CCDC141, SLITRK3, SFRP1, P2RY1, MAP2, VWC2, MAP6, LRP2, SEZ6 |
| regulation of hormone levels | 7 | 15.56 | 4.92E-04 | PRKCA, GCNT4, LRAT, SFRP1, P2RY1, ANO1, APLN |

| | | | | |
|---|---|---|---|---|
| cell development | 13 | 28.89 | 5.88E-04 | DCC, COBL, ACTA1, GABRA5, EPHA3, FAM65B, PRDM14, SLITRK3, SFRP1, MAP2, VWC2, MAP6, SEZ6 |
| dendrite development | 5 | 11.11 | 7.27E-04 | DCC, COBL, MAP2, MAP6, SEZ6 |
| neuron differentiation | 10 | 22.22 | 9.44E-04 | DCC, COBL, SLITRK3, SFRP1, MAP2, GABRA5, VWC2, MAP6, SEZ6, EPHA3 |
| cell-cell signaling | 11 | 24.44 | 1.07E-03 | PRKCA, HECW1, GPR176, CXCL5, SFRP1, P2RY1, ANO1, GABRA5, GRIN2A, SEZ6, APLN |
| locomotion | 11 | 24.44 | 1.14E-03 | PRKCA, DCC, CCDC141, PRDM14, CXCL5, SFRP1, P2RY1, GRIN2A, IGFBP3, EPHA3, IGFBP5 |
| localization of cell | 10 | 22.22 | 1.68E-03 | PRKCA, DCC, CCDC141, PRDM14, CXCL5, SFRP1, P2RY1, IGFBP3, EPHA3, IGFBP5 |
| cell motility | 10 | 22.22 | 1.68E-03 | PRKCA, DCC, CCDC141, PRDM14, CXCL5, SFRP1, P2RY1, IGFBP3, EPHA3, IGFBP5 |
| generation of neurons | 10 | 22.22 | 1.93E-03 | DCC, COBL, SLITRK3, SFRP1, MAP2, GABRA5, VWC2, MAP6, SEZ6, EPHA3 |
| regulation of hormone secretion | 5 | 11.11 | 2.12E-03 | PRKCA, SFRP1, P2RY1, ANO1, APLN |
| cell migration | 9 | 20.00 | 3.22E-03 | PRKCA, DCC, CCDC141, CXCL5, SFRP1, P2RY1, IGFBP3, EPHA3, IGFBP5 |
| hormone secretion | 5 | 11.11 | 3.64E-03 | PRKCA, SFRP1, P2RY1, ANO1, APLN |

**Path 5 Genes:** TNFRSF8, PADI2, AIM1L, FABP3, LCK, LPAR3, TSPAN2, TXNIP, FAM110C, APOB, DPYSL5, ACTG2, TCF7L1, ACOXL, B3GALT1, ITGA6, SLC16A14, SLC6A11, DLEC1, HESX1, CP, VEPH1, PEX5L, MUC4, DNAH5, RASGRF2, POLR3G, KCNN2, DPYSL3, RNF144B, TTBK1, SLC29A1, FILIP1, AIM1, AKAP7, ICA1, PNMA2, VN1R51P, ZNF483, PAPPA, HMCN2, LIPA, SLC16A12, TLL2, JAKMIP3, ADM, BDNF, GYLTL1B, SYTL2, RND1, METTL7A, CHGA, LINGO1, ARRDC4, NECAB2, PIPOX, ARHGAP23, ERBB2, POLR3GP2, PLA2G4C, PTPRT, D21S2088E, MAP7D2, CHST7

| Term | Count | % | PValue | Genes |
|---|---|---|---|---|
| nervous system development | 15 | 23.81 | 6.65E-03 | TSPAN2, SLC6A11, ERBB2, DPYSL5, PADI2, LPAR3, DPYSL3, DNAH5, HESX1, LINGO1, APOB, BDNF, RND1, ADM, TTBK1 |
| cell projection organization | 11 | 17.46 | 7.23E-03 | LINGO1, BDNF, ITGA6, TSPAN2, ADM, FAM110C, ERBB2, DPYSL5, LPAR3, DPYSL3, DNAH5 |
| positive regulation of apoptotic process | 7 | 11.11 | 9.35E-03 | TXNIP, PNMA2, ITGA6, ADM, RASGRF2, LCK, TNFRSF8 |
| positive regulation of programmed cell death | 7 | 11.11 | 9.74E-03 | TXNIP, PNMA2, ITGA6, ADM, RASGRF2, LCK, TNFRSF8 |
| neuron development | 9 | 14.29 | 1.08E-02 | LINGO1, BDNF, RND1, TSPAN2, ADM, ERBB2, DPYSL5, LPAR3, DPYSL3 |
| glycerolipid catabolic process | 3 | 4.76 | 1.10E-02 | APOB, FABP3, PLA2G4C |
| positive regulation of cell death | 7 | 11.11 | 1.23E-02 | TXNIP, PNMA2, ITGA6, ADM, RASGRF2, LCK, TNFRSF8 |
| axon development | 6 | 9.52 | 1.34E-02 | LINGO1, BDNF, TSPAN2, ERBB2, DPYSL5, LPAR3 |
| lipid catabolic process | 5 | 7.94 | 1.42E-02 | APOB, LIPA, ACOXL, FABP3, PLA2G4C |
| neuron projection development | 8 | 12.70 | 1.44E-02 | LINGO1, BDNF, TSPAN2, ADM, ERBB2, DPYSL5, LPAR3, DPYSL3 |
| cellular lipid catabolic process | 4 | 6.35 | 2.02E-02 | APOB, ACOXL, FABP3, PLA2G4C |
| locomotion | 11 | 17.46 | 2.10E-02 | APOB, CHGA, BDNF, ITGA6, FAM110C, ERBB2, LCK, DPYSL5, PADI2, DPYSL3, DNAH5 |
| movement of cell or subcellular component | 12 | 19.05 | 2.39E-02 | APOB, CHGA, BDNF, ITGA6, FAM110C, ERBB2, LCK, KCNN2, DPYSL5, PADI2, DPYSL3, DNAH5 |
| positive regulation of collateral sprouting | 2 | 3.17 | 3.14E-02 | BDNF, LPAR3 |
| leukocyte migration | 5 | 7.94 | 3.15E-02 | APOB, CHGA, ITGA6, LCK, PADI2 |

**Path 6 Genes:** HES3, KLHDC7A, FOXD3, PAX8, GBX2, SORBS2, HTR1A, CTB-180C19.1, CTC-286N12.1, COL12A1, YWHAZP4, VGF, RP11-132A1.3, ARC, ZNF322P1, TRIM22, LINC00678, SIX6, ARHGAP36, SOX3

| Term | Count | % | PValue | Genes |
|---|---|---|---|---|
| regionalization | 4 | 23.53 | 1.89E-03 | ARC, HES3, PAX8, GBX2 |
| midbrain-hindbrain boundary morphogenesis | 2 | 11.76 | 3.12E-03 | HES3, GBX2 |
| pattern specification process | 4 | 23.53 | 4.19E-03 | ARC, HES3, PAX8, GBX2 |
| embryo development | 5 | 29.41 | 5.05E-03 | HES3, PAX8, GBX2, COL12A1, FOXD3 |
| sensory organ development | 4 | 23.53 | 6.39E-03 | SOX3, PAX8, GBX2, SIX6 |
| positive regulation of transcription from RNA polymerase II promoter | 5 | 29.41 | 6.89E-03 | HES3, PAX8, GBX2, SIX6, FOXD3 |
| midbrain-hindbrain boundary development | 2 | 11.76 | 7.01E-03 | HES3, GBX2 |
| embryonic morphogenesis | 4 | 23.53 | 8.87E-03 | HES3, PAX8, GBX2, COL12A1 |
| rostrocaudal neural tube patterning | 2 | 11.76 | 9.33E-03 | HES3, GBX2 |
| transcription from RNA polymerase II promoter | 6 | 35.29 | 9.40E-03 | SOX3, HES3, PAX8, GBX2, SIX6, FOXD3 |
| regulation of transcription from RNA polymerase II promoter | 6 | 35.29 | 9.74E-03 | SOX3, HES3, PAX8, GBX2, SIX6, FOXD3 |
| anterior/posterior pattern specification | 3 | 17.65 | 1.03E-02 | ARC, HES3, GBX2 |
| positive regulation of nucleic acid-templated transcription | 5 | 29.41 | 1.69E-02 | HES3, PAX8, GBX2, SIX6, FOXD3 |
| positive regulation of transcription, DNA-templated | 5 | 29.41 | 1.69E-02 | HES3, PAX8, GBX2, SIX6, FOXD3 |
| positive regulation of RNA biosynthetic process | 5 | 29.41 | 1.78E-02 | HES3, PAX8, GBX2, SIX6, FOXD3 |

**Supplemental Table 2. List of probes used for single-cell qPCR analysis.**

| | | | | | | | |
|---|---|---|---|---|---|---|---|
| ACTB | CHAT | FGF5 | GBX2 | ITGB4 | NANOG | POU4F2 | SLC2A2 |
| ALB | COL10A1 | FOXA1 | GDF3 | KRT10 | NESTIN | POU5F1 | SLC32A1 |
| APLNR | COMP | FOXD3 | GFAP | KRT14 | NEUROD1 | PROM1 | SMTN |
| APOH | CPA1 | FOXG1 | GSC | KRT19 | NEUROG2 | PTCRA | SOX17 |
| AQP1 | CTSK | G6PC | HAND1 | LEFTY1 | NKX2-2 | RCVRN | SOX2 |
| B2M | DCN | GAD1 | HAND2 | MAP3K12 | NKX2-5 | RPLP0 | SOX7 |
| BMP4 | DCX | GAD2 | HES5 | MIOX | NPPA | RUNX1 | T |
| CCR5 | DNMT3B | GALC | HNF4A | MIXL1 | OLIG2 | RYR2 | TAT |
| CD34 | DPP4 | GAPDH | HPRT1 | MSLN | OTX2 | SFTPB | TUBB3 |
| CD3E | ENO1 | GATA1 | IBSP | MYH1 | PAX6 | SFTPD | TYR |
| CD79A | EOMES | GATA2 | IGF2 | MYH7 | PDGFRA | SLC17A6 | ZFP42 |
| CER1 | FABP7 | GATA6 | INS | MYL3 | PODXL | SLC17A7 | ZIC1 |

**Supplemental Table 3. Primer sequences used in qPCR gene expression analysis.**

| Gene | Forward Sequence | Reverse Sequence |
|---|---|---|
| OCT4 | TCAGCCAAACGACCATCTGCCG | AGCAAGGGCCGCAGCTTACA |
| SOX2 | TACAGCATGTCCTACTCGCAG | GAGGAAGAGGTAACCACAGGG |
| Nanog | ACGCAGAAGGCCTCAGCACCTA | AGGTTCCCAGTCGGGTTCACCA |
| T | ACCTGTGTCGCCACCTTCCA | ACCACTGGCTGCCACGACAA |
| MIXL1 | TCCTCAACCACTGTGCTCCTGG | AACCCCGTTTGGTTCGGGCA |
| EOMES | AGGCGCAAATAACAACAACACC | ATTCAAGTCCTCCACGCCATC |
| GSC | CGCGGGACACTTGCCCGTATTA | AAGGCAGCGCGTGTGCAAGA |
| PAX6 | CCAGAAAGGATGCCTCATAAA | TCTGCGCGCCCCTAGTTA |
| Nestin | CCGCATCCCGTCAGCTGGAAAA | GCTTGGGCACAAAAGCCAGCA |
| OTX2 | CTTAAGCAACCGCCTTACGC | AGGGGTGCAGCAAGTCCATA |
| AFP | AGCTGACCTCGTCGGAGCTGAT | TCCCTCGCCACAGGCCAATAGT |
| KDR | ACCGTTAAGCGGGCCAATGGA | ACCACGGCCAAGAGGCTTACCT |
| GAPDH | TTCTTTTGCGTCGCCAGCCG | TGACCAGGCGCCCAATACGA |
| EF1a | GCTGGCTTCACTGCTCAGGTGATT | TGCAATGTGAGCCGTGTGGCA |

### Supplemental Experimental Procedures

#### Spatial Analysis

Spatial analysis for cells seeded as colonies at different split ratios or single-cells at different densities was performed using the SpatStat package for R (Baddeley and Turner, 2005; Baddeley et al., 2015). The spatial coordinates of each cell within a field (i.e. each microscopic image) were converted into a spatial point pattern, and each field was divided into 5x5 grids (totaling 25 quadrats). The number of points (or cells) per quadrat was quantified using the 'quadratcount' feature in Spatstat for each quadrat in each image acquired for a given well. The Coefficient of Variation (CV; standard deviation divided by mean) was then calculated for the number of cells per quadrat for each seeding condition. For each well, the total number of cells was quantified, and normalization of CV values was performed by calculating the CV of a simulated random uniform distribution of points (equal to the cell number in that well) using the 'runifpoint' function in Spatstat.

#### Bioinformatics

FASTQ sequencing data for each sample was aligned to GENCODE version 23 human genome annotations (hg38), using the HISAT2 alignment tool (version 2.0.1) (Kim et al., 2015). The 'featureCounts' function from the Rsubread package (version 1.22.3) (Liao et al., 2019)was used to count reads per gene, which were passed onto DESeq2 (version 1.12.4) (Love et al., 2014)for library size normalization and detection of differentially expressed genes (FDR <= 0.05). Hierarchical clustering was done with the R 'heatmap.2' function from the gplots package (version 2.17.0) for differentially expressed genes. Principal component analysis was done using an in-house R script and the built-in R PCA function 'prcomp'. Temporal expression path clustering was generated by unsupervised hierarchical clustering a log2 fold (log2FC) change matrix using R's built-in 'hclust' function. This was done for all genes with an absolute log2FC of at least 2 in any single timepoint. Clusters which were visibly similar were then manually combined, and cluster trajectories were then plotted using an in-house R script. Gene set enrichment analyses (Subramanian et al., 2005) were conducted by comparing differentially expressed genes from RNA-seq data to custom Gene Sets produced in-house. The -log10(p-value) for each gene from RNA-seq data was used as a custom weighting when GSEA analysis was conducted (version 2.1.0). Gene ontology analysis was performed using DAVID version 6.8 (Huang et al., 2009) and the BINGO plugin for Cytoscape (Maere et al., 2005).

#### Single-cell Gene Expression Analysis

Cycle numbers to enrichment for each gene in each cell were normalized to housekeeping genes between cells. Normalized cycle numbers were converted to a cycle difference measurement, subtracting observed cycle number from the mean cycle number of the corresponding gene across all cells. An aggregate vector of all cycle differences was used to produce Z-scores for each gene in each cell; these Z-scores were then used to plot heatmaps, using the R 'heatmap.2' function from the gplots package (version 2.17.0). t-SNE plots were also produced from cycle differences using the Rtsne package (version 0.10) and an in-house script for plotting the results. Tissue and cell type enrichment for single cell expression data was done using Enrichr (Chen et al., 2013; Kuleshov et al., 2016).

#### Chondrocyte Micromass Differentiation and Analysis

Following 48-hour pre-differentiation in E8, E6 or BA, cells (H9 hESCs or iPSCs) were dissociated using TrypLE Express and resuspended in chondrogenic media supplemented with 10 μM Y-27632. Chondrogenic media consisted of high glucose DMEM (Life Technologies) containing 1% KOSR, 1% ITS+ premix (BD Biosciences), 1% Sodium Pyruvate, 1% non-essential amino acids, 1% Penicillin-Streptomycin (Life Technologies), 100 μg/mL ascorbic acid 2-phosphate, $10^{-7}$ M dexamethasone, and 40 μg/mL L-proline (Sigma). Cells were resuspended at a density of $2x10^7$ cells/mL and plated as 15 μL drops in Matrigel-coated 12-well plates for 2 hours, after which 1 mL of chondrogenic media containing Y-27632 was added. Fresh chondrogenic media was added daily for course of the 7-day protocol.

To assess differentiation by matrix production, 4-5 micromass cultures were pooled and digested overnight at 65°C using 40 μg/mL papain enzyme in digestion buffer and processed for quantification as described (Lee et al., 2011). Sulfated glycosaminoglycan (s-GAG) content was quantified using Dimethylmethylene blue (DMMB). Hydroxyproline (OH-Pro) content was quantified by acid hydrolysis of papain-digested samples followed by neutralization, oxidation, and addition of 4-dimethylaminobenaldehyde. Hoechst 33258 dye was used to quantify DNA content in the samples, and total s-GAG or OH-Pro content were normalized to amount of DNA. Proteoglycan production was assessed after 7 days by Alcian Blue staining. Micromass cultures were fixed in 4% PFA for 15 min, rinsed with 0.2N HCl, and stained with 0.1% Alcian Blue solution at pH1 (diluted with 0.2N HCl from 0.3% w/v Alcian blue in 70% ethanol solution; Sigma) overnight at room temperature. Cultures were rinsed thoroughly with distilled water and imaged using a Zeiss Axio Zoom V16.

#### Endothelial Progenitor Cell Differentiation and Analysis

Differentiation of iPSCs derived from late-EPCs (described in Chang et al., 2013) was performed using an adapted protocol from Tatsumi *et al* (Tatsumi et al., 2011). Briefly, cells were pretreated for 48 hours with E6 supplemented with 10 μg/ml of both BMP4 and Activin A. Subsequently, cells were cultured in DMEM/F12 (Life Technologies) supplemented with B27 (Life Technologies), N2 (Life Technologies) and BIO (Sigma) for 72 hours with daily media change performed. Cells were then cultured in StemPro-34 (Life Technologies) supplemented with 50 μg/ml of $VEGF_{165}$ (R&D Systems) for a further 48 hours prior to MAC-selection with CD144

microbeads (Miltenyi Biotech). CD144 enriched cells were then further expanded and routinely passaged in StemPro-34 media supplemented with 50 μg/ml $VEGF_{165}$.
Endothelial cell differentiation was assessed by flow cytometry against a panel of endothelial cell surface markers. Briefly, following six days of differentiation, cells were harvested and dissociated into single cell suspension using TrypLE. A total of 2 $x10^5$ cells in 200 μl of flow buffer was enumerated and aliquotted into each tube. Directly conjugated antibodies (all from BD Bioscience) to CD31 (Cat. No. 340297), CD34 (Cat. No. 345802), CD45 (Cat. No. 555482), CD144 (Cat. No. 560411) and VEGFR (Cat. No. 560494) were then added at the recommended manufacturer's dilution into each tube (i.e. 20 μl per test) and incubated on ice in the dark for 30 mins. To halt the staining process, 400 μl of flow buffer was added per tube and the entire volume of 600 μl was subjected to centrifugation at 400rpm for 5 mins. Supernatant was decanted and cells resuspended in 300 μl flow buffer prior to analysis on an Attune acoustic focus cytometer (Life Technologies). To gate samples for FACS analysis, cells (20,000 events) were initially gated by FSC-A vs SCS for the exclusion of debris. Data were analyzed using FlowJo v10.0.6 software.

**Primary Antibodies**

OCT4A (Cell Signaling Technology C52G3; 1:400), Brachyury T (R&D Systems AF2085, 5 μg/mL), SOX2 (R&D Systems MAB2018, 8 μg/mL; EMD Millipore AB5603, 1:200), Nestin (EMD Millipore ABD69, 1:500), Desmin (Abcam ab191181, 1:200) and SOX17 (R&D Systems AF1974, 1:100).

**Supplemental References**


Baddeley, A., and Turner, R. (2005). **spatstat** : An *R* Package for Analyzing Spatial Point Patterns. J. Stat. Softw.

Baddeley, A., Rubak, E., and Turner, R. (2015). Spatial Point Patterns: Methodology and Applications with {R} (London: Chapman and Hall/CRC).

Chang, W.Y., Lavoie, J.R., Kwon, S.Y., Chen, Z., Manias, J.L., Behbahani, J., Ling, V., Kandel, R.A., Stewart, D.J., and Stanford, W.L. (2013). Feeder-independent derivation of induced-pluripotent stem cells from peripheral blood endothelial progenitor cells. Stem Cell Res. *10*, 195–202.

Chen, E.Y., Tan, C.M., Kou, Y., Duan, Q., Wang, Z., Meirelles, G., Clark, N.R., and Ma'ayan, A. (2013). Enrichr: interactive and collaborative HTML5 gene list enrichment analysis tool. BMC Bioinformatics *14*, 128.

Huang, D.W., Sherman, B.T., and Lempicki, R.A. (2009). Systematic and integrative analysis of large gene lists using DAVID bioinformatics resources. Nat. Protoc. *4*, 44–57.

Kim, D., Langmead, B., and Salzberg, S.L. (2015). HISAT: a fast spliced aligner with low memory requirements. Nat. Methods *12*, 357–360.

Kuleshov, M. V, Jones, M.R., Rouillard, A.D., Fernandez, N.F., Duan, Q., Wang, Z., Koplev, S., Jenkins, S.L., Jagodnik, K.M., Lachmann, A., et al. (2016). Enrichr: a comprehensive gene set enrichment analysis web server 2016 update. Nucleic Acids Res. *44*, W90-7.

Lee, W.D., Hurtig, M.B., Kandel, R.A., and Stanford, W.L. (2011). Membrane Culture of Bone Marrow Stromal Cells Yields Better Tissue Than Pellet Culture for Engineering Cartilage-Bone Substitute Biphasic Constructs in a Two-Step Process. Tissue Eng. Part C Methods *17*, 939–948.

Liao, Y., Smyth, G.K., and Shi, W. (2019). The R package Rsubread is easier, faster, cheaper and better for alignment and quantification of RNA sequencing reads. Nucleic Acids Res.

Love, M.I., Huber, W., and Anders, S. (2014). Moderated estimation of fold change and dispersion for RNA-seq data with DESeq2. Genome Biol. *15*, 550.

Maere, S., Heymans, K., and Kuiper, M. (2005). BiNGO: a Cytoscape plugin to assess overrepresentation of gene ontology categories in biological networks. Bioinformatics *21*, 3448–3449.

Subramanian, A., Tamayo, P., Mootha, V.K., Mukherjee, S., Ebert, B.L., Gillette, M.A., Paulovich, A., Pomeroy, S.L., Golub, T.R., Lander, E.S., et al. (2005). Gene set enrichment analysis: a knowledge-based approach for interpreting genome-wide expression profiles. Proc. Natl. Acad. Sci. U. S. A. *102*, 15545–15550.

Tatsumi, R., Suzuki, Y., Sumi, T., Sone, M., Suemori, H., and Nakatsuji, N. (2011). Simple and highly efficient method for production of endothelial cells from human embryonic stem cells. Cell Transplant. *20*, 1423–1430.
